# Newly Sequenced Genomes Reveal Patterns of Gene Family Expansion in Select Dragonflies (Odonata: Anisoptera)

**DOI:** 10.1101/2023.12.11.569651

**Authors:** Ethan R. Tolman, Christopher D. Beatty, Paul B. Frandsen, Jonas Bush, Or R. Bruchim, Ella Simone Driever, Kathleen M. Harding, Dick Jordan, Manpreet K. Kohli, Jiwoo Park, Seojun Park, Kelly Reyes, Mira Rosario, Jisong L. Ryu, Vincent Wade, Anton Suvorov, Jessica L. Ware

**Author notes:** Corresponding author; 2085139703; 829 Canyon Ridge Rd #303, Blacksburg, VA 24060.

## Abstract

Gene family evolution plays a key role in shaping patterns of biodiversity across the tree of life. In Insecta, adaptive gene family turnover has broadly been tied to vision, diet, pesticide resistance, immune response and survival in extreme environments. Patterns of gene family evolution are of particular interest in Odonata (dragonflies and damselflies), which represents the first lineage to fly, and one of the most exceptional groups of predators. Previous work in Odonata found expansions of opsin genes are correlated with the diversification of the herbivorous insects that Odonata prey upon, but general trends in gene family turnover has not been studied in this order. Here, we show that two families of suborder Anisoptera (dragonflies), Libellulidae and Petaluridae, have expanded gene repertoire related to their unique life history and diversification patterns. These results are an important step towards understanding why Libellulidae is, generally, a species rich family of short lived species that are highly tolerant to poor water quality, while Petaluridae is a species poor family of habitat and behavioral specialists. Specifically, Libellulidae share expanded gene families related to immune response, desiccation response, and processing of free radicals, which all potentially enable many Libellulidae to inhabit low quality water bodies. Likewise, Petaluridae show unique patterns of gene turnover in gene families implicated in sensory perception, which could be tied to the unique semi-terrestrial lifestyle of the nymphs of this family. Furthermore, Odonata as a whole has a gene gene turnover rate that is an order of magnitude smaller than other studied insect orders, potentially contributing to the relatively low species diversity in the order Odonata compared to other insects. These results offer important hypotheses for the consideration of evolutionary drivers across Insecta.

## Introduction

Gene turnover, through both duplication and loss, is a crucially important source of both variation and novel adaptation across the tree of life (Innan and Kondrashov 2010). In particular, rapidly evolving gene families, with high divergence between related species, are more closely related to adaptation than other genes in the genome (Zhang et al. 2013; Kim et al., 2022). While insects exhibit a stunning array morphological, physiological and behavioral adaptations are thought to have influenced their success-flight, herbivory and complete metamorphosis are all suggested to be dramatic drivers of insect diversity (Misof et al. 2014)-the implication of gene turnover in these adaptations is not well studied across all insect taxa, with a particular lack of taxonomic coverage in aquatic insect lineages (Hotaling et al. 2020). A better understanding of adaptive gene family evolution may help explain why insects have been so successful (Luo et al., 2018); work on insect genomes has already revealed how turnover in gene families is related to adaptations of species and whole lineages.

While not meant to be exhaustive, the following are pertinent hypotheses of adaptive gene family evolution in insects. Diversification of supergene families is a major force behind pesticide resistance in mosquitoes (Ranson et al. 2002; Claret et al. 2024). The expansion of gene families associated with detoxification can be related to polyphagy in the Lepidoptera family Noctuidae (Breeschoten et al., 2022), and in species of Thysanoptera (Hu et al., 2023). Detoxifying genes have also duplicated in the weevil *Rhynchophorus ferrugineus* (Olivier, 1790), as have genes related to insecticide resistance (Hazzouri et al., 2020). Expansion of gene families related to nutrient metabolism, development, stress response, reproduction, and immune processes may play a major role in the adaptability of the fall armyworm, *Spodoptera frugiperda* (Smith, 1797; Gao et al., 2023). The expansion of sensory related gene families can be associated with host differences in Hymenoptera (Kuang et al., 2023). The Antarctic midge *Parochlus stenenii* (Gercke, 1889) has undergone many gene family expansions tied to processes such as the innate immune system, unfolded protein response, DNA packaging, protein folding, and unsaturated fatty acid biosynthesis that are likely adaptive to the frigid environment it inhabits (Kim et al., 2022). The expansion of transcription factors, proteases, odorant receptors, and horizontally transferred genes involved in nutritional symbioses probably play an important role in interactions with plants and bacteria in the hemipteran *Bactericera cockerelli* (Šulc, 1909; Kwak et al., 2023). Gene family expansion has also been linked to adaptations necessary for life in the ocean in the family Hermatobatidae (Wang et al., 2023). Duplications of genes within the cytochrome p450 family, and families of olfactory binding proteins are associated with pollution tolerance and mate finding in the caddisfly S*tenopsyche tienmushanensis* (Hwang, 1957; Luo et al., 2018). Opsins genes are highly diverse, with numerous expansions and contractions across the insect tree of life, but the evolutionary processes driving these patterns largely remain unclear (Guignard et al., 2022).

In dragonflies, gene family evolution could provide insight into the question: why have some taxonomic lineages diversified into many species while others remained species poor? The most diverse family in Anisoptera, Libellulidae, likely originated in the late Paleogene (∼20-40Ma; Suvorov et al., 2021) and has since radiated into over 1500 known species. Libellulidae inhabit a wide variety of aquatic habitats, and some taxa are very tolerant of poor water quality (Šigutová et al., 2022). Although generation times have not been studied for most of the Libellulidae, they are generally understood to have, on average, a relatively rapid life-cycle of less than one year, compared to other families of Odonata (Tennessen, 2019).

In contrast to Libellulidae, the least diverse family within Anisoptera is Petaluridae, which originated in the mid-to-late Jurassic (∼150-175 Ma; Ware et al., 2014a; Kohli et al., 2021; Suvorov et al., 2021; Tolman et al., 2024), and contains only 11 extant species, with little fossil evidence of widespread extinction (Ware et al., 2014a; Tolman et al., 2024). The ages of the individual species lineages within Petaluridae range between 15 and 115 million years (Ware et al., 2014a; Tolman et al., 2024). While Libellulidae inhabit a broad range of habitats, Petaluridae only inhabit fens with high water quality (Turner, 1970; Baird, 2012; Ware et al., 2014a; Baird, 2019). The lifespan of Petaluridae varies from 5 to 20 years depending on the species, quite possibly making them the longest lived dragonflies (Baird, 2012).

With the exception of micro-chromosomes, the genomes of these two families differ very little on a structural scale, as the genomes of the petalurid *Tanypteryx hageni* (Selys, 1879) and libellulid *Pantala flavescens* (Fabricius, 1798) share widespread synteny both with each other, and with two distantly related species of Zygoptera (Tolman, Beatty, et al., 2023).

Within this framework of remarkable karyotypic stability, there is some evidence that gene family evolution is tied to phenotypic and behavioral diversity in Odonata. Opsin duplication and losses have been correlated with both larval and adult behavior, and occurred around the diversification of flowering plants, and herbivorous insects which Odonata prey upon (Futahashi et al., 2015; Suvorov et al., 2017). Despite promising initial data from opsins, patterns of gene family evolution more broadly have not been considered in Odonata. Identifying gene family expansions can illuminate the molecular processes behind the phenotypic and ecological diversity of Odonata and should lead to further research into the functional genomics of these Odonata, which may be tied to adaptations to their different habitats. An understanding of patterns of gene family evolution also informs how species may respond to shifting habitats during climate change (Kim et al., 2022). Similarly, comparative work exploring different levels of gene turnover across the Odonata tree of life, can shed light upon the diversity of different clades (Thomas et al., 2020).

To better understand the patterns of gene family evolution in the dragonfly families Libellulidae and Petaluridae, and determine how gene expansion is related to the evolution and ecology of the two families, we sequenced and annotated the genomes of the libellulid *Pachydiplax longipennis* (Burmeister, 1839) and the petalurid *Uropetala carovei* (White, 1846). We then included the annotations in a gene family expansion analysis of all publicly available genomes of Odonata (two Libellulidae, one Petaluridae, and three Zygoptera) that were generated with PacBio HiFi data at the time of analysis. Because it is relevant for understanding gene family expansion, we also briefly discuss the life history of each of the five Anisoptera used in the analysis below.

### Pachydiplax longipennis (Libellulidae)

Perhaps no dragonfly interacts with humans more than the monotypic species *Pachydiplax longipennis*. Within its range, which spans the United States, Southern Canada, Mexico, and several Caribbean Islands, *P. longipennis* is the most observed aquatic insect, and fortieth most observed species on the community science platform iNaturalist, despite being one of the least observed Anisoptera in more conserved habitats such as National Parks, suggesting this species may predominantly be utilizing urban habitats (Tolman et al. 2024b). Indeed *P. longipennis* was the most commonly observed Odonata by community scientists in a recent survey of the Odonata in a North American City (Uche-Dike et al. 2024). Further supporting the hypothesis that *P. longipennis* thrives in urban habitats, some genes involved in the response to oxidative stress, immune response, DNA damage repair, all functions implicated in urban adaptation (Salmón et al. 2018), were suggested to be under diversifying selection while other genes involved in these processes are under stabilizing selection (Tolman et al., 2024b). The range of *P. longipennis* recently has expanded northward with a warming climate (Lis et al., 2020). Reflecting its seeming propensity for urban habitats, *P. longipennis* can be observed in localities with a highly diverse assemblage of Odonata, as well as bodies of water with few species, suggesting a high tolerance to poor water quality (Tollett et al., 2009). Many cohorts of *P. longipennis* develop over the course of a year, but bi-voltinism has also been observed in this species (Wissinger, 1988).

### Pantala flavescens (Libellulidae)

*P. flavescens,* the wandering glider, is one of two species in *Pantala.* It is among the most well-studied species of Anisoptera, and is the most widely distributed dragonfly (Russell et al., 1998). It makes long transcontinental and trans-oceanic migrations (Rowe, 2004; Ware et al., 2022), and is frequently found in warmer regions, particularly in the tropics, but can also be found in temperate regions in the summer (May, 2013). Though it is not surprising—given the migratory nature of the species—evidence suggests widespread gene flow between populations across the globe, and it may be appropriate to refer to many, if not all, of the migratory sub-populations as panmictic (Ware et al., 2022). An analysis of historical effective population size (N_e_), using individuals from Guangdong Province, China found that the N_e_ of *P. flavescens* has varied between 1 million and 10 million over the last million years, with declines associated with the onset of the agricultural and industrial period (H. Liu et al., 2022).

### Sympetrum striolatum (Charpentier, 1840; Libellulidae)

*S. striolatum* is a palearctic dragonfly (John Horne, 2012) that is found in Europe and other areas in the Mediterranean (Borkenstein and Jödicke, 2022), with bodies of salt-water likely serving as a major barrier to gene flow within this species (Parkes et al., 2009). A demographic model of *S. striolatum*, generated from a reference assembly of an individual from Wytham Woods, Oxfordshire, UK (Crowley et al., 2023) estimated a relatively stable N_e_ around 1 million throughout most of the past 10 million years (which may predate the divergence times of this species), with declines to an effective population size of around 100,000 occurring in the last 100,000 years (Tolman, Bruchim, et al., 2023).

### Tanypteryx hageni (Petaluridae)

*T. hageni* is one of only two species in *Tanypteryx* and is the only Petaluridae to inhabit the fens of Western North America, ranging from Southern California, USA, to Southern British Columbia, Canada (Turner, 1970). *T. hageni* burrows in its nymphal stage, and takes five years to emerge as an adult (Turner, 1970). A recent behavioral analysis demonstrated that *T. hageni* nymphs regularly leave their burrows for extended periods of time during both day and night, navigating a terrestrial environment. *Tanypteryx* is one of two genera in the “Laurasian” clade of Petaluridae (Tolman et al., 2024)

### Uropetala carovei (Petaluridae)

*U. carovei* is one of two species of the genus *Uropetala* (referred to by Māori as Kapokapowai), and is found in the lowlands of both the Northern and Southern islands of New Zealand (Rowe, 1981). Their lifespan ranges from five to six years, depending upon the length of the growing seasons (L. S. Wolfe, 1952). Like *T. hageni*, *U. carovei* burrows in its nymphal stage (Green, 1977). It has been suggested that *U. carovei* leaves its burrow to forage (G. Theischinger and I. Endersby, 2009), but it is unclear if *U. carovei* is as active outside of its burrow as *T. hageni*. *Uropetala* (Kapokapowai) is one of three genera in the “Gondwanan” clade of Petaluridae.

## Materials and Methods

### Specimen collection

We sequenced the genome of *U. carovei*, a member of the Gondwanan clade of Petaluridae which diverged from the Laurasian clade of Petaluridae (including *T. hageni*) in the late Jurassic (Ware et al., 2014a; Tolman et al., 2024). *U. carovei* was collected as a nymph near Aorere goldfields in New Zealand and was injected with and stored in 100% ethanol prior to DNA extraction and sequencing. To expand the taxon coverage of the Libellulidae we sequenced the genome of *Pachydiplax longipennis*. An adult specimen was collected from Parkcenter Park Pond in Boise, Idaho, and was stored in 100% ethanol prior to sequencing.

### Sequencing and Genome Size Estimation

High molecular weight DNA was extracted from both the *U. carovei* and *P. longipennis* specimens using the Qiagen Genomic-tip kit. High molecular weight DNA was sheared to 18 kbp using a Diagenode Megaruptor and separate libraries were prepared for each specimen using the PacBio HiFi SMRTbell Express Template Kit 2.0. Each library was sequenced on a single PacBio Revio SMRT cell in CCS mode at the BYU DNA sequencing center.

### Genome Assembly and QC

For both *U. carovei* and *P. longipennis* we used HiFi reads to estimate genome size with Genomescope 2.0 (Ranallo-Benavidez et al., 2020). We first generated read histograms with a histogram length of 21 using Jellyfish (Marçais and Kingsford, 2011), and ran Genomescope 2.0 with default settings. We generated both whole genome assemblies with hifiasm v.0.16.0 (Cheng et al., 2021).

We used taxon-annotated GC-coverage plots generated with BlobTools v1.1.1 (Laetsch and Blaxter, 2017) to screen for contamination. We mapped all HiFi reads against the final assembly for each species using minimap2 v2.1 (Li, 2018) and sorted the resulting bam files with samtools v1.13 (Danecek et al., 2021). We then assigned taxonomy with megablast (Shiryev et al., 2007) using the parameters: task megablast and -e-value 1e-25. We calculated coverage using the blobtools function map2cov, created the blobtools database for each species using the command blobdb, and generated the blobplot with the command “`blobtools plot`”. After examining the blobplots we removed one contig from the assembly of *P. longipennis* which was assigned to the phylum tracheophyta and removed all contigs from the assembly of *U. carovei* with coverage above 100x, or below 10x, which did not blast to Arthropoda.

Additional contaminants were automatically removed from the assembly of *U. carovei* as a part of the NCBI genome upload (supplementary Supplementary Table 1). No additional contaminants were removed from the assembly of *P. longipennis*.

We generated all contiguity stats for both assemblies with the command line tool assembly-stats v0.1.4 (Trizna, 2020). We assessed the completeness of both assemblies with BUSCO v.4.1.4 (Manni et al., 2021) using the Insecta Odb10 database, in genome mode with the flag -–long to retrain the initial BUSCO results for a more accurate identification of BUSCO genes.

### Whole genome alignment

Given the contiguity of the assembly of *P. longipennis*, we calculated synteny between the draft genome assembly and the chromosome length genome assembly of *S. striolatum* to identify any complete chromosomes. We identified potentially homologous genes using BLASTp (Camacho et al., 2009), restricting output to 5 hits per gene and filtering for hits with an e-value below 1e-5. We combined the BLAST output and gff annotation files for *P. longipennis* and *S. striolatum* and used MCscanX (Wang et al., 2012) to identify areas of synteny between the genomes with a minimum of 5 genes. We visualized the syntenic relationships in a Circos v.0.69-18 (Krzywinski et al., 2009) plot to identify chromosome-length contigs.

To assemble the remaining contigs into chromosomes, we aligned the chromosome length assemblies of *Pantala flavescens*, *Sympetrum striolatum*, *Tanypteryx hageni* and the Platycnemididae *Platycnemis pennipes* with the assembly of *P. longipennis* using progressive cactus v.2.6.7 (Armstrong et al., 2020). We then used ragout v.2.3 (Kolmogorov et al., 2014; Kolmogorov et al., 2018), with the option --solid-scaffolds, to generate a scaffolded assembly from the alignment. As the karyotype of *U. carovei* is currently unclear (Kuznetsova and Golub, 2020), it would not be possible to assess the quality of the scaffolding, so we did not attempt to scaffold this genome. This would not impact the results of the gene family evolution analysis.

### Gene family expansion analysis

#### Data selection

In addition to the genomes of *P. longipennis* and *U. carovei* we included publicly available genomes of Odonata from GenBank (October 1, 2023). To minimize bias due to differing genome sequencing methods we only included genomes that had been generated from PacBio high-fidelity technology, which has been shown to generate the highest quality insect genomes to date (Hotaling et al., 2021). We identified four Anisopteran genomes sequenced with this technology, comprising two Libellulidae (*Pantala flavescens* (H. Liu et al., 2022) and *Sympetrum striolatum* (Crowley et al., 2023)) and one Petaluridae (*Tanypteryx hageni* (Tolman et al., 2023)). We also chose to include the three publicly available genome assemblies of Zygoptera, the Platycnemididae *Platycnemis pennipes* (Benjamin W. Price and E. Louise Allan, 2023), the Coenagrionidae *Ischnura elegans* (Price et al., 2022), and the Calopterygidae *Hetaerina americana* (Grether et al., 2023) to use as an outgroup in the gene family expansion analysis.

#### Annotation

To ensure consistency in the genome annotations we annotated with genomes of *H. americana*, *P. pennipes, I. elegans, P. longipennis, S. striolatum, U. carovei,* and *T. hageni* in GALBA (Brůna et al., 2023) using the annotated protein set of *P. flavescens* (H. Liu et al., 2022). GALBA aligns the reference protein set to the soft-masked target genome using miniprot (Li, 2023) and uses the alignments to train Augustus (Stanke et al., 2006; Hoff and Stanke, 2019), and has produced more accurate annotations of insect genomes than common annotation pipelines such as BRAKER2 (Brůna et al., 2021; Brůna et al., 2023) which use RNA-seq. Of the published genomes of Odonata, the annotated protein set of *P. flavescens* (H. Liu et al., 2022) is by far the most complete (Tolman et al., 2023), with nearly 99% of BUSCO genes present (fig 1.A), justifying its use as a reference. Prior to annotation with GALBA, repetitive elements of all target species were modeled and soft-masked with RepeatModeler2 (v2.0.1) and RepeatMasker v.4.1.2-p1(Flynn et al., 2020). Because the initial GALBA annotations found an unreasonably high (>50,000) number of genes, we only kept annotated genes with a significant BLAST (Camacho et al., 2009) hit (with an e-value cutoff of 1e-10) to the annotated protein set of *P. flavescens* (H. Liu et al., 2022). Given the completeness of this annotated protein set (H. Liu et al., 2022) this filtering step increases the specificity of GALBA without a drop in sensitivity, and has since been incorporated into the GALBA pipeline (see https://github.com/Gaius-Augustus/GALBA/releases/tag/v1.0.10). The GALBA documentation indicates this filtering step removes erroneous annotations, and with a low homology threshold, retains accurate gene models. The documentation contains detailed analyses demonstrating that such a filtering step is highly unlikely to remove accurate gene models, and is thus unlikely to miss key orthologues and bias a downstream gene family expansion analysis. The new filtered annotations are much more complete than the original genome annotations for the species (Tolman et al., 2023). We generated all contiguity stats for both assemblies with the command line tool assembly-stats v0.1.4 (Trizna, 2020). We assessed the completeness of all annotations with BUSCO v4.1.4 (Manni et al., 2021) using the Insecta ODB10 database, in protein mode.

**Figure 1:**
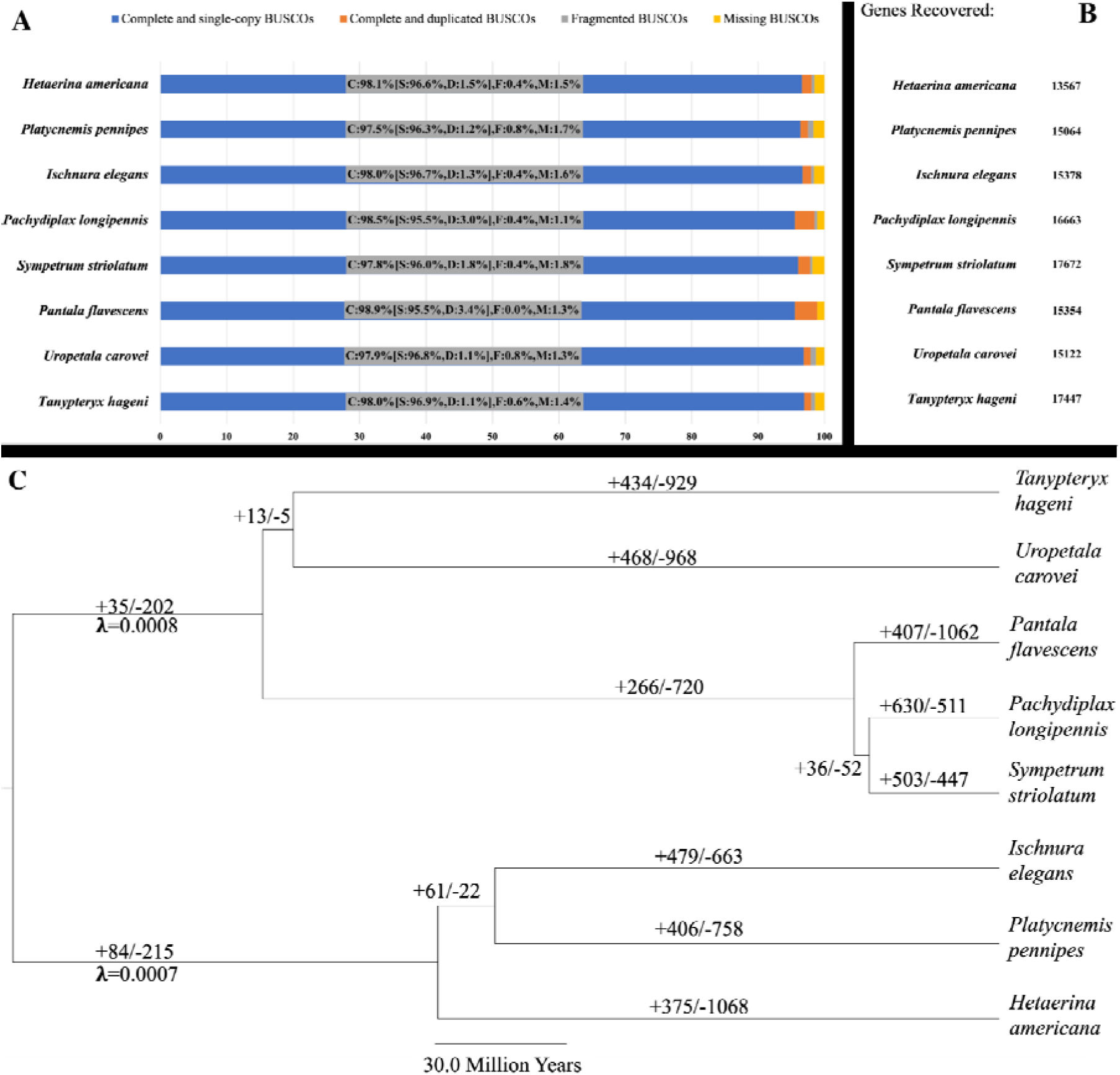
(A) BUSCO results of the eight utilized Odonata genomes. (B) Number of genes recovered with GALBA (Brůna et al., 2023) annotation, after filtering out genes that did not have a BLAST match with the genome annotation of *P. flavescens.* (C) Results of global lambda gene family expansion analysis. The global lambda is shown at the root with the number of identified expansions and contractions on each branch. *Annotation previously reported (H. Liu et al., 2022)

As part of the annotation process, we also identified the mitochondrial genomes from the de novo assembles of *P. longipennis* and *U. carovei* using mitohifi v. 3.0.0 (Uliano-Silva et al., 2021). We used the mitochondrial genome of *T. hageni* (Tolman et al., 2023) as a reference for *U. carovei* and the mitochondrial genome of *S. striolatum* (Crowley et al., 2023) as a reference for *P. longipennis*. The mitochondrial genome sequences have been uploaded, as part of the reference genome sequences, to GenBank.

#### Gene family evolution

For the ultrametric input tree required by CAFE5 (Mendes et al., 2021), we largely used divergence times recovered by Suvorov et al. (2021) which utilized transcriptomic datasets, and shared overlapping interfamilial divergence times with Kohli et al. (2021). The dated tree from Suvorov et al. (2021) contained Platycnemididae, while the tree by Kohli et al. (2021) did not, making it the more prudent of the two time-trees to utilize here. The only branch length not used from this analysis was the divergence between the Petaluridae *T. hageni* and *U. carovei,* which was set to 161 Ma, as suggested by Tolman et al. (2024), which used a multilocus anchored-hybrid enrichment dataset. Suvorov recovered a more recent split (∼115 Ma) between the Gondwonan (including *Uropetala*) and Laurasian (including *Tanypteryx*) clades in Petaluridae by placing the fossil *Argentinopetala archangelski* (F. Petrulevičius and Nel 2003) as crown Petaluridae. Tolman et al. (2024) demonstrated that, based on wing venation, this fossil should be considered as a crown member of the Gondwanan clade of Petaluridae. This fossil placement altered the divergence time estimation between the two clades within Petaluridae, but did not alter the divergence estimation between Petaluridae and other families in Odonata, which closely overlapped with the time identified by Suvorov et al. (2021) (Tolman et al., 2024). While *Pachydiplax longipennis* and *Platycnemis pennipes* were not included in the dated tree considered by Suvorov et al. (2021), this tree does include other taxa which shared the same most-recent common ancestor of *Sympetrum striolatum* and *Pachydiplax longipennis* (the genus *Libellula* is more closely related to *Pachydiplax* than is *Sympetrum* (Ware et al., 2007)) and *Ischnura elegans* and *Platycnemis pennipes* (other representative of Platycnemididae were included by Suvorov et al.) so these dates were used in our ultrametric tree.

We clustered the amino acid sequences from the longest isoforms of each gene for each species with Orthofinder (Emms and Kelly, 2019). We then used CAFE5 (Mendes et al., 2021) to generate three different lambda (gene birth/death rate) models of gene family evolution. First, a global lambda model with a shared lambda for the whole tree, second a two lambda model with separate lambdas for Anisoptera and Zygoptera, and third a four lambda model with separate lambdas for Petaluridae, Libellulidae, the ancestral branch of Anisoptera, and Zygoptera. We ran each model with 100,000,000 iterations to allow for convergence and replicated each model run 1000 times, which involved first running CAFE5 with the ultrametric tree and orthofinder output to generate an error model, and then rerunning CAFE5 with the estimated error model to account for sequencing and annotation error for each of the replicate runs. We then identified significantly expanding and contracting gene families from each converged model and used the Akaike Information Criterion (AIC) and Likelihood-ratio Test (LRT) to compare each of the converged models. In addition to comparing the converged models with AIC and LRT, we also randomly selected 1000 combinations of each of the model runs without resampling, and identified the number of times each model was selected by AIC and LRT in the random comparison. Finally, to assess convergence we evaluated the variance of -LnL values estimated across each of the 1000 runs for each lambda model. Specifically we tested whether the -LnL standard deviation (SD) of the two and four lambda models significantly differ from the SD of the global model using the F-test in R v4.1.4 (R Core Team 2021) to determine whether the independent runs of the two and four lambda runs varied more than the global lambda run.

While the four lambda model did outperform the other two models in AIC and LRT and was the best fitting model by both metrics when comparing model runs independently (Supplementary Table 3), its -LNL SD of this model indicated it did not converge as strongly as the other two models (Supplementary Table 3), with a particular lack of convergence around the lambda for the ancestral branch of Anisoptera (supplementary Fig. 6). This lack of convergence is likely the cause of the differing counts of expansions and contractions in the four lambda model as compared to the other two tested models (Supplementary Table 4).The likelihood surface may have a flat plateau near the optimum or have a multiple local optima near the optimum Due to this lack of convergence, we considered less complex models for the the summary of the functions of expanded gene families, as the CAFE5 documentation recommends use convergence as a strong consideration in considering the appropriateness of gene family evolution models (hahnlab/CAFE5 2025). In selecting between the global and two lambda models, we chose to use expanded gene families selected from the two lambda model, as it provides better fit than the global lambda model in the AIC and LRT tests, was selected at a higher rate (11.3% overall) than the global model (0.3% overall) by both AIC and LRT, and showed small improvement in the strength of convergence compared to the global model (Supplementary Table 3). Additionally, these two models had 93% overlap in the identification of significantly expanding gene families, and only minor differences in the counts of expanding and contracting gene families (Supplementary Table 4).

#### Functional annotation

We obtained Gene Ontology (GO) terms for all of the protein sequences (with the exception of *P. flavescens* which were previously reported (H. Liu et al., 2022) using interproscan (Mariani et al., 2013; Baek et al., 2021; Hiranuma et al., 2021). We summarized the GO terms of significantly expanding gene families into TreeMaps with REVIGO (Supek et al., 2011) for expanded gene families in the Libellulidae and Petaluridae. Given the missing taxon sampling in other areas of the tree, it would not be appropriate to analyze patterns of gene family expansion on a broader level.

#### Demographic histories

As the demographic history of a species may be important when studying genome evolution, particularly in relict insect lineages lineages (H.-L. Liu et al., 2022; Bentley et al., 2023), we estimated the demographic histories of *U. carovei*, *P. longipennis* and *T. hageni* (estimates for all other species have previously been described (H. Liu et al., 2022; Tolman, Bruchim, et al., 2023)). As with the blobtools analysis, HiFi reads were mapped to the genome assemblies with minimap2 v2.1.1(Li, 2018), and the bam files were sorted with samtools v1.16.1 (Danecek et al., 2021) and bcftools v1.6 (Danecek et al., 2021), with a minimum depth of 5 and a maximum depth of 100, was used to make base calls from the sorted bam files. We then used psmc v0.6.5-r67 (Liu and Hansen, 2017) to convert the fastq basecalling output to psmcfa format and estimate the pairwise sequential markovian coalescent (PSMC) with 100 bootstrap replicates for each species. For all three species we visualized the PSMC with the programs psmc (Liu and Hansen, 2017) and gnuplot v5.2 (Phillips, 2012) using four different mutation rates (1e-9, 2e-9, 3e-9, and 4e-9), encompassing known genome-wide mutation rates in Insecta (Liu et al., 2017), and values previously used in demographic analyses of Odonata (H. Liu et al., 2022; Ethan R. Tolman et al., 2023). As *P. longipennis* cohorts are typically uni-voltine, but occasionally exhibit bi-voltinism (Wissinger, 1988), we used a generation time of 0.8 to reflect the occasional bi-voltine generations. *U. carovei* has a generation time of 5-6 years depending on the length of growth seasons (Winstanley and Rowe, 1980; Baird, 2012) so we used a generation time of 5.5 years. *T. hageni* has a life cycle of 5 years (Turner, 1970; Baird, 2012), and this was the value used in the coalescent analysis.

## Results

### Sequencing and Genome Size Estimation

We generated over 6 million high-fidelity reads, totaling more than 93 Gbp, for *U. carovei* (read N50=15,101; L50=2,488,241; ave=14,547.95) and over 5.8 million reads totaling more than 76 Gbp for *P. longipennis* (read N50 = 13,394; L50 = 2,369,807; ave= 13,034.64). Using GenomeScope 2.0, we estimated a genome size of 0.88 Gb (87.4% unique sequence, 1.8% heterozygosity, kcov: 40.5, err: 0.0885%, k:21) for *P. longipennis* (supplementary fig. 1A). and 1.00 Gb (67.5% unique sequence, 1.18% heterozygosity, kcov: 36.7, err: 0.224%, k:21) for *U. carovei* (supplementary fig. 1B).

### Genome Assembly and QC

The initial assembly of *U. carovei* was highly contiguous when compared to other whole genome assemblies of Odonata (length=1.33Gb, N50=8.67Mb, L50=49, ave=1.72Mb, N=773). The assembly of *P. longipennis* was among the most contiguous genomes ever assembled without scaffolding (length=1.0Gb, N50=66.9 1Mb, L50=7, ave=3.71Mb, N=270). The blobplot, which displays the assigned taxonomy, coverage and GC content of each contig to identify potential contaminants, of *U. carovei* showed potential contamination (supplementary fig. 2A) of contigs with coverage above 100x, or below 10x. Such contigs that were not assigned to Arthropoda were removed. Only one contig of *P. longipennis*, which was assigned by BLAST to tracheophyta (a common laboratory plant) was removed from the assembly (supplementary fig. 2B), as all other contigs were assigned to Arthropoda. The identification of core BUSCO orthologues indicate that the genomes were highly complete, with nearly 97 percent of assessed Insecta genes present in the assembly of *U. carovei* (C:96.9%[S:95.9%,D:1.0%],F:1.0%,M:2.1%,n:1367), and 98% of the Insecta genes present in *P. longipennis* (C:98.0% [S:95.2%,D:2.8%], F:0.5%, M:1.5%, n:1367). Both are comparable to other HiFi assemblies of Odonata (Newton et al., 2023 Jun 7; Tolman et al., 2023).

### Whole genome alignment

We generated eight pseudo-chromosomes for the assembly of *P. longipennis* from the cactus whole genome alignment with *S. striolatum* (supplementary Table 1). The remaining four chromosomes still showed high levels of synteny with a select number of contigs (supplementary Table 1, supplementary fig. 3).

### Annotation

All our annotations recovered more than 97% of BUSCO Insecta db10 genes (fig. 1A) after filtering. We initially recovered 114,332 genes for *H. americana*, 103,410 for *P. pennipes*, 66,745 for *I. elegans*, 45,043 for *P. longipennis*, 53,217 for *S. striolatum*, 53,629 for *U. carovei*, and 75,637 for *T. hageni*. After filtering for genes that had a significant BLAST hit with the proteome of *P. flavescens* we recovered 13,567 genes for *H. americana*, 15,064 from *P. pennipes*, 15,378 from *I. elegans*, 16,663 from *P. longipennis*, 17,672 from *S. striolatum*, 15,122 from *U. carovei*, and 17,447 from *T. hageni* (fig. 1B).

### Gene family evolution

#### Gene birth-death rate

The four lambda model had a lower AIC (258702) than the global (261966) and two (261974) lambda models, and both the two lambda (94) and four lambda models outperformed the Global lambda in the LRT (Supplementary Table 3). However, the four lambda model did not converge as well as the global lambda model, with a significantly higher standard deviation of -LnL from 1000 (SD=1872.92) independent runs than the global model (SD=170.67; Supplementary Table 3). The two lambda model had a significantly smaller distribution of final model -LnL than the global model (SD=132.67; Supplementary Table 3).

The gene families identified as significantly evolving by CAFE shared 93% overlap between the one and two lambda models, 88% overlap between the one and four lambda models, and 83% overlap between the two and four lambda models. The counts of identified expanding and contracting gene families differed by a max of 13 (in Anisoptera) between the one and two lambda models (Supplementary Table 4). The four lambda model did not identify any expansions or contractions in the ancestral Anisoptera, and the branches leading to Petaluridae, Libellulidae, and Zygoptera differed from the global model by counts of 11, 72, and 157 respectively. We identified convergence at □ = 0.0008 in the global lambda model (Fig. 1C; supplementary Fig. 4), □ = 0.0008 and □ = 0.0007 for Anisoptera and Zygoptera respectively in the two lambda model (supplementary Fig. 5), and □ = 0.0000091(ancestral Anisoptera), □ = 0.00058 (Petaluridae), □ = 0.002 (Libellulidae), and □ = 0.007 (Zygoptera) in the four lambda model (supplementary Fig. 6).

#### Functional annotation

GO terms associated with significantly expanding gene families shared amongst the Libellulidae and Petaluridae in the two lambda model were summarized into 44 (Fig. 2A) and 3 (Fig 2B) biological processes respectively. Within Libellulidae, GO terms associated with gene family expansion were summarized into 44 biological processes in *S. striolatum*, 13 in *P. flavescens*, and 45 processes in *P. longipennis* (Fig. 2A). In Petaluridae, GO terms from significantly expanded gene families were summarized into 36 biological processes in *T. hageni*, and 32 processes in *U. carovei*.

**Figure 2.**
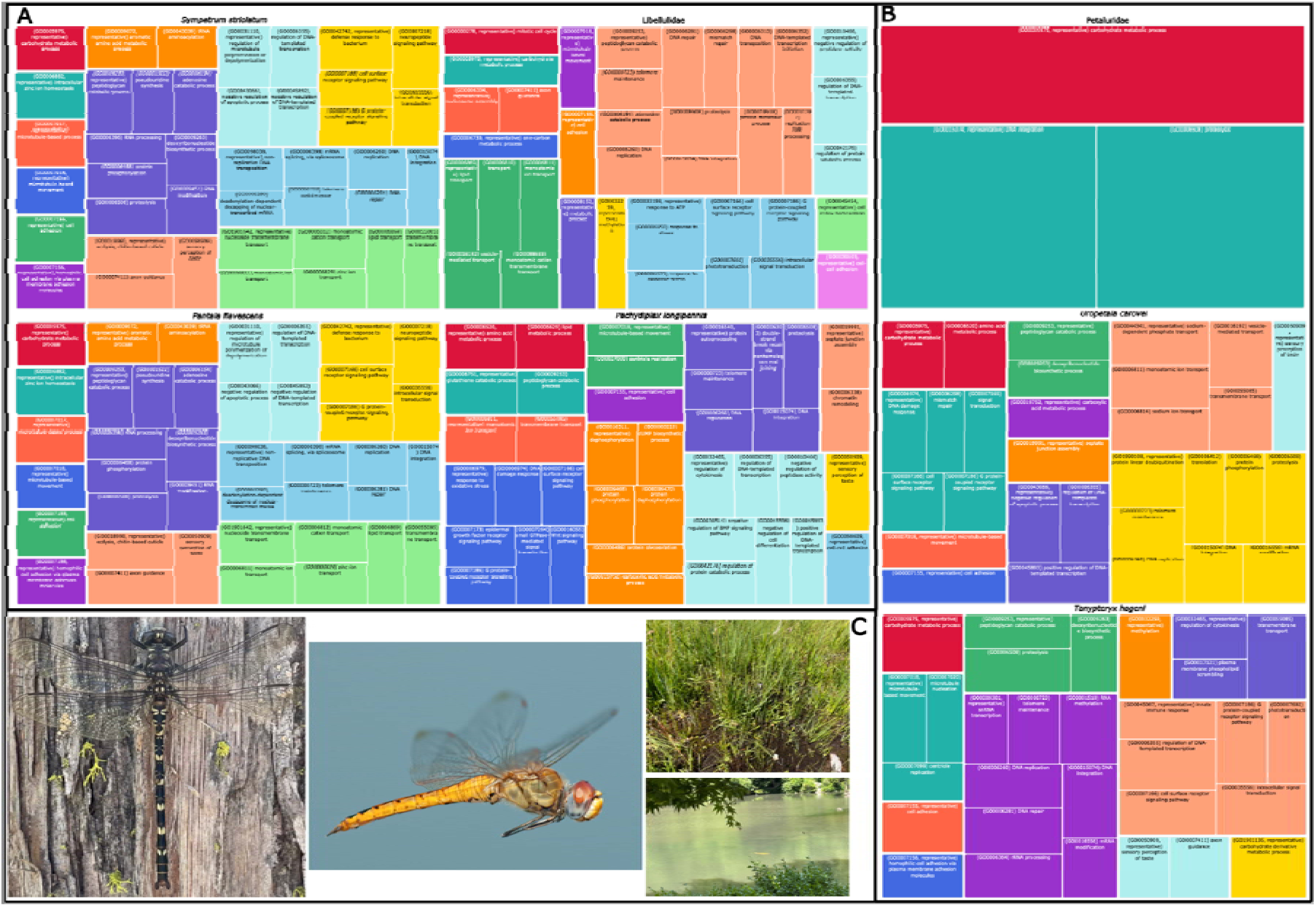
TreeMaps summarizing rapidly expanding gene families in the (A) Libellulidae and (B) Petaluridae. Shared gene family expansions are represented under “Libellulidae” and “Petaluridae”. The enriched biological process terms were summarized with REVIGO (Supek et al., 2011). (C) From left to right, an adult of the petalurid Tanypteryx hageni, an adult of the libellulid Pantala flavescens, (top) a fen habitat inhabited by T. hageni (California, USA) and (bottom) an urban pond where the authors have observed both P. longipennis and P. flavescens (New York, USA)

To test the assumption that the Petaluridae have long been habitat specialists, and the Libellulidae have long been generalists, we modeled the historical effective population size of the species considered here, as habitat specialists are more likely to have a historically smaller effective population size. PSMC models showed one climb in N_e_ of *P. longipennis*, starting between 7 and 20 Mya, and peaking from 1-6 Mya (fig. 3A). The peak N_e_ was estimated to be between 0.5 and 6 million, with declines stabilizing less than 100k years before present at ∼100,000 Fig. 3A). The PSMC model suggests the N_e_ of *Uropetala carovei* fluctuated throughout a period from 25-100 Mya depending on the mutation rate (fig. 3B). In the scenario with the lowest genome-wide mutation rate, from 100 Mya to 10 Mya, the N_e_ of *Uropetala* increases, but continuously decreases until around a million years ago. In the scenario with the fastest mutation rate, the population starts increasing around 25 Mya and peaks around 2.5 Mya (fig. 3B). Among the different scenarios the highest peak N_e_ was between 7 and 25 million (fig. 3B).

**Figure 3:**
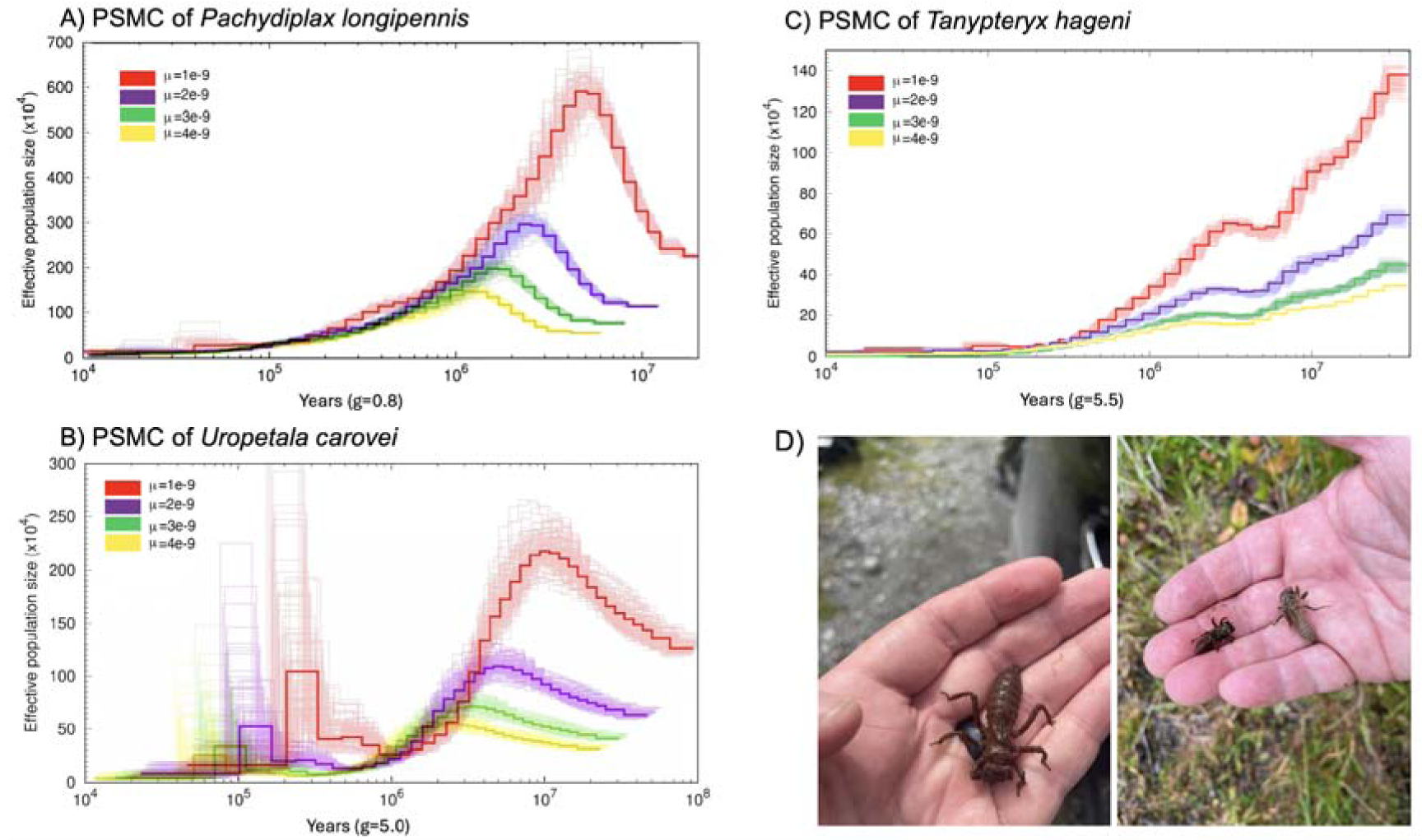
Demographic histories of A) P. longipennis, B) U. carovei, and C) Tanypteryx hageni estimated with the Pairwise Sequential Markovian Coalescent (PSMC) using mutations of 1,2,3, and 4e-9. Bootstrap replicates are shown as lighter lines. A generation time of .8 years was used for P. longipennis, 5 years for T. hageni and 5.5 years for U. carovei. D) The semi-terrestrial nymphs of U. carovei (left) and T. hageni (right) in their specialized fen habitats.

All estimates for the change in effective population size (N_e_) for *T. hageni* showed an initial drop in N_e_ starting around 30 million years ago (Mya), until a brief uptick in N_e_ approximately 3 Mya, followed by another period of decline stabilizing approximately 200,000 years before present (fig. 3C), as made evident by the tight bootstrap replicates. Regardless of the mutation rate used in the model, the N_e_ of *T. hageni* never climbed higher than 2 million, and lower estimates of the genome-wide mutation rate do not show an effective population size above 1 million (fig. 3C).

## Discussion

Our results provide a number of testable hypotheses relating to genome size, gene family expansion, and the diversification of odonates into the range of ecosystems they currently inhabit. We consider here the strengths and implications of the technologies used to generate these results, and then some of the major research areas suggested by our results.

### Genome assembly and annotation

Genome size estimates of Odonata assembled from PacBio high-fidelity sequences have been found to diverge from genome sizes estimated with Genomescope (Ranallo-Benavidez et al., 2020). Although many contigs from the *U. carovei* assembly were not assigned to the Arthropoda, it is likely an artifact of an incomplete NCBI reference database (Tolman et al., 2023). Another Petaluridae, *T. hageni*, showed a similar profile with contigs not assigned to Arthropoda, even in well supported scaffolds with HiC data, likely due to poor coverage in NCBI databases (Tolman et al., 2023). This phenomenon has also been observed in horseshoe crabs, and in the transcriptomes of other Petaluridae (Tolman et al., 2023). With our current understanding, taxonomically mis-assigned contigs with a similar GC content and coverage to contigs not assigned to Arthropoda are not a major source of concern (Tolman et al., 2023).

PacBio high-fidelity sequencing has become the gold standard for insect genomics (Hotaling et al., 2021). The near chromosome-level assembly of *P. longipennis* presented here, before any scaffolding, further demonstrates the potential of this sequencing technology. As this technology is further developed, complete, chromosome-length genome assemblies will be available without any scaffolding.

The genome annotation tool GALBA has been demonstrated to outperform more established pipelines, such as BRAKER2 (Brůna et al., 2021) when annotating insect genomes, and does not need further sequencing of transcriptomes (Brůna et al., 2023). While our genome annotations initially had a high (>50,000) number of genes compared to the annotation of *P. flavescens*, by conservatively retaining only genes which had a BLAST hit to a protein sequence from *P. flavescens* (H. Liu et al., 2022), we were able to generate highly complete annotations with nearly all insecta BUSCO orthologues present (Fig. 1. A, B), further supporting the efficacy of this pipeline in insects, especially in the absence of transcriptomic data.

### Gene turnover in Odonata

The global lambda recovered for Odonata (□=0.0008) was higher than a previous estimate of 0.000010 (Thomas et al., 2020). However this previous estimate only included one species in Odonata, which could explain this disparity. Our estimate is still much lower than other animal lineages, as the estimated lambdas of primates (□=0.00563), birds of paradise (□=0.00226), and the insect orders Hymenoptera (□=0.00375) (Mendes et al., 2021), and Hemiptera (□= 0.00233070.01; Kwak et al., 2023) are all an order of magnitude greater than the global lambda of Odonata (□=0.0008). An analysis of the synteny between chromosome level genome assemblies of Odonata found a high degree of conservation in chromosome structure, even between suborders (Tolman, Beatty, et al., 2023). Thus, our results here demonstrate that not only does the karyotype of Odonata evolve much slower than many studied insect lineages, but gene turnover in Odonata also appears to occur at a much slower rate than other studied insect lineages. Such a dynamic could partially explain why the Odonata are so species poor (with ∼6000 species) compared to other insect lineages.

It is also notable that that, in the four lambda model, the rate of gene turnover in the Petaluridae (□ = 0.00058) was an order of magnitude lower than the gene turnover rate identified in Libellulidae (□ = 0.002). This low turnover rate could partially explain why Petaluridae has not radiated to the extent of Libellulidae (nor is there evidence of widespread extinction in the Petaluridae (Tolman et al., 2024)). While this is certainly an exciting hypothesis for future study, caution does need to be taken at this time, as the lambda for ancestral anisoptera did not show strong evidence of convergence (Supplementary Fig 6; Table 1). This hypothesis should be further explored as more Odonata genomes become available, and as bioinformatic methods for studying gene family evolution improve.

### Hypotheses from expanding gene families

While more research is necessary to better understand the molecular processes tied to the evolutionary and ecological histories of odonates, our functional annotations of expanding gene families can be used to generate testable hypotheses related to the ecology and evolution of Petaluridae and Libellulidae. Here, we discuss hypotheses derived from our GO analyses that warrant further investigation in future studies.

As *T. hageni* and *U. carovei* represent ∼18% of all Petaluridae, and both major clades within the family (Ware et al., 2014, Tolman et al., 2024), we believe it is highly likely that most of the gene family expansions shared between these two species are shared across Petaluridae. However, as the family Gomphidae is likely sister to Petaluridae (Bybee et al., 2021; Kohli et al., 2021; Suvorov et al., 2021), without representative genomes for Gomphidae, we cannot determine whether these expanded gene families would be found in both families, thereby suggesting that they may have originated in a most-recent common ancestor. Given these limitations, the expanding gene families shared between the two Petaluridae still suggest candidate genes to explore for genomic adaptations related to the life history attributes of this family, namely the longevity of these lineages through evolutionary time, the long development times seen in these species, and the unique semi-terrestrial lifestyle of the nymphs.

#### Hypothesis 1: terrestrial nymphs have different sensory perception profiles

The genomes of *T. hageni* and *U. carovei* both contained expanded gene families implicated in the “sensory perception of taste.”(fig. 2B). While all odonate nymphs are aquatic, the nymphs of this petaltail species have been observed to forage in the terrestrial habitats around their subterranean burrows (which fill with water, providing an aquatic environment). Odonates are generalist predators in their nymph and adult stages. Terrestrial foraging could bring these petalurid nymphs in contact with a range of potential prey very different from those they would encounter in a strictly aquatic environment. Insects generally utilize taste to avoid toxic chemicals, find suitable food sources, select mates and find oviposition sites (Rebora et al., 2022). Adult Odonata, in particular, have been observed catching and rejecting toxic prey upon tasting it (Rebora et al., 2022). It is currently unclear if nymphs exhibit this behavior. With terrestrial foraging, petalurid nymphs may encounter a greater proportion of toxic prey, such as arachnids,which have been observed in and around the burrows of *T. hageni*. Further examination of the specific genes involved in taste perception in petalurids needs to be performed, along with greater documentation of their foraging behavior and analysis of their diet to test this hypothesis.

Our results also suggested that *T. hageni* perceives light differently than other studied Anisoptera. The expansion of gene families related to *phototransduction* in *T. hageni* could be due to the differing light conditions the nymphs face in a terrestrial environment. Transcriptomic data has shown that opsin genes in particular have undergone a number of expansions and contractions across Odonata, but the reasons for this evolutionary pattern are currently unclear (Bybee et al., 2022). Transcriptomic work will be needed to confirm whether expanded gene families related to phototransduction are expressed in the nymphs, as previously published transcriptomic work including Petaluridae sequenced only adults (Suvorov et al., 2017). It is currently unclear if *U. carovei* is semi-terrestrial to the same extent as *T. hageni*, and it is necessary to determine the extent of semi-terrestriality across Petaluridae to refine any hypotheses related to this behavior. As the nymphal behavior of the Petaluridae is better understood, and more genomes and transcriptomes are sequenced, gene families associated with taste and vision could be a promising avenue for understanding their interaction with a terrestrial environment as nymphs.

#### Hypothesis 2: highly tolerant dragonflies have expanded gene families to deal with desiccation and bacterial exposure

While it is certainly plausible that shared expansions in the three Libellulidae analyzed here are shared across the family, we will be more certain about gene family expansions among libellulids when more genomes are available. Several biological processes related to the shared expansions of these three taxa may be associated with the generalist life-history of libellulids. A major overrepresented GO term shared between these three species is the *peptidoglycan catabolic process* (fig. 2A). Peptidoglycan is an essential element in the walls of most bacteria (Vollmer et al., 2008). Several hypotheses could explain why gene families related to catabolizing peptidoglycan would be present. First: genes involved in the peptidoglycan catabolic process were upregulated when the Antarctic midge *Belgica antarctica* was placed under desiccation conditions (Teets et al., 2012). In the case of odonates, expansions could be adaptive to deal with more arid environments, or in ephemeral aquatic environments. Secondly, Libellulidae often inhabit warm, eutrophic waters, where harmful bacteria are more likely to be found (Sharip and Fauzi, 2019). Expansion of gene families involved in the peptidoglycan catabolic process could enable Libellulidae to tolerate exposure to high levels of bacteria.

#### Hypothesis 3: highly tolerant dragonflies have expanded gene families to deal with exposure to free radicals

Another GO term associated with expanded gene families in Libellulidae is *cell redox homeostasis* (fig. 2A), a dynamic but essential process ensuring balance between oxidizing and reducing reactions in cells that is also involved in many biological responses (Le Gal et al., 2021). These elements have been shown to maintain homeostasis in fluid conditions (Le Gal et al., 2021). An ability to maintain homeostasis would be critical for Libellulidae, as the shallow bodies of water they inhabit can often undergo rapid changes in temperature and water chemistry. GO terms associated with expanding gene families of *P. longipennis* could offer some explanation for how this species has been so successful in colonizing urban environments. One of the most overrepresented GO terms for this species was *response to oxidative stress* (fig. 2A). Birds in urban or polluted environments have increased antioxidant protection, and face higher levels of oxidative stress (Salmón et al., 2018), so it is likely that Odonata face these same pressures. It follows that a greater capacity to respond to oxidative stress in urban environments would allow *P. longipennis* to outcompete other species that do not have this capacity. Additionally, the expansion of gene families related to *chromatin remodeling* (fig. 2A) could allow *P. longipennis* sufficient plasticity to thrive in an urban environment.

#### Hypothesis 4: highly tolerant dragonflies have expanded gene families to allow for greater phenotypic plasticity

Expansions related to both *methylation*, *response to ATP*, and *response to stress* in Libellulidae (fig. 2A) could result in increased phenotypic and molecular plasticity, which are a necessity for evolutionary stability in fluctuating environments (Lalejini et al., 2021), and could be another molecular adaptation to fluctuating environments.

#### Hypothesis 5: migratory dragonflies can more efficiently process carbohydrates

In *P. flavescens*, the expansion of gene families related to the *carbohydrate metabolic process* are a plausible mechanism for allowing this species to undertake trans-oceanic and trans-continental migrations in a single generation (Ware et al., 2022). It is likely that an especially efficient process for obtaining energy from carbohydrates would be selected for on such long migrations. The genome of closely related non-migratory species to *P. flavescens* are needed to explore the genomic adaptations to a migratory lifestyle.

We recognize that a limitation of this study is a lack of taxonomic coverage. *T. hageni* and *U. carovei* represent the two major clades of Petaluridae (Laurasian and Gondwanan) and ∼18% of the family. However, the three species of Libellulidae anlayzed here represent only 3 of 29 genera and 3 of >1500 species. Additionally, our analysis includes only 2 of the 39 families of Anisoptera, 3 of the 39 Zygoptera, and only 8 of ∼6300-6400 species of Odonata (May, 2019; Waller et al., 2019), so we acknowledge the birth/death rates calculated could be influenced by poor taxon sampling, and more genomes are needed to more accurately model gene birth and death. We have included all of the high-quality odonate genome assemblies currently available at the time of analysis, offering predictions about birth/death rates in this order that can be refined as more data becomes available.

### Demographic histories

Similar to *I. elegans* and *S. striolatum* (Tolman, Bruchim, et al., 2023) the PSMC model for *P. longipennis* showed a decline in effective population size (N)_e_ roughly corresponding with the Pleistocene ice ages. As the climate warms this species could become more widely distributed, but more research into the ecological niche of this species is needed to affirm this assumption. The N_e_ of the two fen-dwelling petalurid species–*U. carovei* and *T. hageni*–were estimated to be smaller than *P. longipennis* or the other libellulids through time (H. Liu et al., 2022; Tolman, Bruchim, et al., 2023). If the Petaluridae have consistently been habitat specialists through the period covered by our analysis, we would predict that their population size would be limited by availability of their habitat.

## Conclusions

Our findings are the first comprehensive analysis of gene family expansion in Odonata. We identify a number of gene families that may be associated with the differing evolutionary and ecological histories of the Petaluridae and the Libellulidae. The role of these gene families, and the relatively low levels of gene turnover will be an exciting avenue of future research as more genomes from across the Odonata, along with more transcriptome data, become available.

## Author contributions

E.R.T: Methodology, formal analysis, data curation, writing.

C.D.B: Methodology, formal analysis, writing.

P.B.F: Methodology, writing, resources.

R.J.B: Methodology, writing.

O.R.B: Formal analysis, writing.

E.S.D: Formal analysis.

K.M.H: Formal analysis.

D.J: Methodology, resources, writing.

M.K.K Methodology.

J.P: Formal analysis.

S.P: Formal analysis.

K.R: Formal analysis, writing.

M.R: Formal analysis.

J.L.R: Formal analysis.

V.W: Formal analysis, writing.

A.S. Methodology, resources, supervision, writing.

J.L.W: Methodology, formal analysis, resources, supervision, writing.

## Acknowledgments

We would like to thank Kristin Gnojewski and Cindy Busche for their support of this project. We acknowledge funding from the Conservation Connection Foundation, the Society of Systematic Biologists Graduate Student Research Award, the 2023 Sydney Anderson Travel Award from the American Museum of Natural History, and the National Science Foundation (DEB#2002473). We thank the people of Aotearoa (New Zealand) for permission to work on their lands and collect specimens of *Uropetala carovei* (DOC Authorization #103630-RES).

## Declaration of interests

The authors declare no competing interests.

## Data availability

The genome assemblies and rawreads of U. carovei (PRJNA1043152 and P. longipennis (PRJNA999334) are available on GenBank. RepeatMasker output (10.6084/m9.figshare.24714909), unfiltered annotations (10.6084/m9.figshare.24714918) and filtered proteins sets used for gene family expansion analysis (10.6084/m9.figshare.24593130) are all available on figshare.

## Notes

### Competing Interest Statement

The authors have declared no competing interest.

### Summary of Updates

Updated to reflect revised gene turnover analysis.

## References

Armstrong J, Hickey G, Diekhans M, et al. 2020. Progressive Cactus is a multiple-genome aligner for the thousand-genome era. Nature. 587(7833):246–251. 10.1038/s41586-020-2871-y.

Baek M, DiMaio F, Anishchenko I, et al. 2021. Accurate prediction of protein structures and interactions using a three-track neural network. Science. 373(6557):871–876. 10.1126/science.abj8754.

Baird I. 2012. The wetland habitats, biogeography and population dynamics of Petalura gigantea (Odonata: Petaluridae) in the Blue Mountains of New South Wales [PhD]. The University of Western Sydney.

Baird IRC. 2019. Establishment of larval pits by Tachopteryx thoreyi (Odonata: Petaluridae): habitat modification by a non-burrowing petalurid. International Journal of Odonatology. 22(2):135–146. 10.1080/13887890.2019.1636889.

Benjamin W. Price, E. Louise Allan. 2023. The genome sequence of the White-legged damselfly, Platycnemis pennipes (Pallas, 1771). Available from https://wellcomeopenresearch.org/articles/8-320.

Bentley BP, Carrasco-Valenzuela T, Ramos EKS, et al. 2023. Divergent sensory and immune gene evolution in sea turtles with contrasting demographic and life histories. Proceedings of the National Academy of Sciences. 120(7):e2201076120. 10.1073/pnas.2201076120.

Borkenstein A, Jödicke R. 2022. Thermoregulatory behaviour of Sympetrum striolatum at low temperatures with special reference to the role of direct sunlight (Odonata: Libellulidae). 51:83–109.

Breeschoten T, van der Linden CFH, Ros VID, et al. 2022. Expanding the Menu: Are Polyphagy and Gene Family Expansions Linked across Lepidoptera? Genome Biology and Evolution. 14(1):evab283. 10.1093/gbe/evab283.

Brůna T, Hoff KJ, Lomsadze A, et al. 2021. BRAKER2: automatic eukaryotic genome annotation with GeneMark-EP+ and AUGUSTUS supported by a protein database. NAR Genomics and Bioinformatics. 3(1):lqaa108. 10.1093/nargab/lqaa108.

Brůna T, Li H, Guhlin J, et al. 2023. Galba: genome annotation with miniprot and AUGUSTUS. BMC Bioinformatics. 24(1):327. 10.1186/s12859-023-05449-z.

Bybee SM, Futahashi R, Renoult JP, et al. 2022. Transcriptomic insights into Odonata ecology and evolution. In: Cordoba-Aguilar A, Beatty C, Bried J, editors. Dragonflies and Damselflies: Model Organisms for Ecological and Evolutionary Research. Oxford University Press. p. 0. 10.1093/oso/9780192898623.003.0003.

Bybee SM, Kalkman VJ, Erickson RJ, et al. 2021. Phylogeny and classification of Odonata using targeted genomics. Molecular Phylogenetics and Evolution. 160:107115. 10.1016/j.ympev.2021.107115.

CAFE. 2023. Available from https://github.com/hahnlab/CAFE5.

Camacho C, Coulouris G, Avagyan V, et al. 2009. BLAST+: architecture and applications. BMC Bioinformatics. 10(1):421. 10.1186/1471-2105-10-421.

Cheng H, Concepcion GT, Feng X, et al. 2021. Haplotype-resolved de novo assembly using phased assembly graphs with hifiasm. Nat Methods. 18(2):170–175. 10.1038/s41592-020-01056-5.

Claret J-L, Di-Liegro M, Namias A, et al. 2024. Despite structural identity, ace-1 heterogenous duplication resistance alleles are quite diverse in Anopheles mosquitoes. Heredity. 132(4):179–191. 10.1038/s41437-024-00670-9

Christopher D. Beatty, Aaron Goodman, Ethan R. Tolman, et al. In Preparation. The Larval Behavior of Tanypteryx Hageni.

Crowley LM, Price BW, Allan EL, et al. 2023. The genome sequence of the Common Darter, Sympetrum striolatum (Charpentier, 1840). Wellcome Open Res. 8:389. 10.12688/wellcomeopenres.19937.1.

Danecek P, Bonfield JK, Liddle J, et al. 2021. Twelve years of SAMtools and BCFtools. Gigascience. 10(2):giab008. 10.1093/gigascience/giab008.

Emms DM, Kelly S. 2019. OrthoFinder: phylogenetic orthology inference for comparative genomics. Genome Biology. 20(1):238. 10.1186/s13059-019-1832-y.

Flynn JM, Hubley R, Goubert C, et al. 2020. RepeatModeler2 for automated genomic discovery of transposable element families. Proceedings of the National Academy of Sciences. 117(17):9451–9457. 10.1073/pnas.1921046117.

G. Theischinger, I. Endersby. 2009. Identification Guide to the Australian Odonata. Hurtsville, New South Wales, Australia: Department of Environment, Climate Change and Water NSW.

Gao H, Li Y, Tian Y, et al. 2023. Gene family expansion analysis and identification of the histone family in Spodoptera frugiperda. Comparative Biochemistry and Physiology Part D: Genomics and Proteomics. 48:101142. 10.1016/j.cbd.2023.101142.

Green LFB. 1977. Aspects of the respiratory and excretory physiology of the nymph of Uropetala carovei (Odonata: Petaluridae). New Zealand Journal of Zoology. 4(1):39–43. 10.1080/03014223.1977.9517934.

Grether GF, Beninde J, Beraut E, et al. 2023. Reference genome for the American rubyspot damselfly, Hetaerina americana. Journal of Heredity. 114(4):385–394. 10.1093/jhered/esad031.

Guignard Q, Allison JD, Slippers B. 2022. The evolution of insect visual opsin genes with specific consideration of the influence of ocelli and life history traits. BMC Ecology and Evolution. 22(1):2. 10.1186/s12862-022-01960-8.

[Software] hahnlab/CAFE5. 2025. [accessed 2025 Mar 25]. https://github.com/hahnlab/CAFE5.

Hazzouri KM, Sudalaimuthuasari N, Kundu B, et al. 2020. The genome of pest Rhynchophorus ferrugineus reveals gene families important at the plant-beetle interface. Commun Biol. 3(1):1–14. 10.1038/s42003-020-1060-8.

Hiranuma N, Park H, Baek M, et al. 2021. Improved protein structure refinement guided by deep learning based accuracy estimation. Nat Commun. 12(1):1340. 10.1038/s41467-021-21511-x.

Hoff KJ, Stanke M. 2019. Predicting Genes in Single Genomes with AUGUSTUS. Curr Protoc Bioinformatics. 65(1):e57. 10.1002/cpbi.57.

Hotaling S, Sproul JS, Heckenhauer J, et al. 2021. Long Reads Are Revolutionizing 20 Years of Insect Genome Sequencing. Genome Biology and Evolution. 13(8):evab138. 10.1093/gbe/evab138.

Hotaling S, Kelley JL, Frandsen PB. 2020. Aquatic Insects Are Dramatically Underrepresented in Genomic Research. Insects. 11(9):601. 10.3390/insects11090601

Hu Q-L, Ye Z-X, Zhuo J-C, et al. 2023. A chromosome-level genome assembly of Stenchaetothrips biformis and comparative genomic analysis highlights distinct host adaptations among thrips. Commun Biol. 6(1):1–10. 10.1038/s42003-023-05187-1.

Innan H, Kondrashov F. 2010. The evolution of gene duplications: classifying and distinguishing between models. Nat Rev Genet. 11(2):97–108. 10.1038/nrg2689

John Horne. 2012. Emergence, maturation time and oviposition in the Common Darter Sympetrum striolatum (Charpentier). Journal of the British Dragonfly Society. 28(2). Available from https://british-dragonflies.org.uk/wp-content/uploads/2020/11/JBDS_Vol28_2.pdf#page=13.

Kim H, Kim H-W, Lee JH, et al. 2022. Gene family expansions in Antarctic winged midge as a strategy for adaptation to cold environments. Sci Rep. 12(1):18263. 10.1038/s41598-022-23268-9.

Kohli M, Letsch H, Greve C, et al. 2021. Evolutionary history and divergence times of Odonata (dragonflies and damselflies) revealed through transcriptomics. iScience. 24(11):103324. 10.1016/j.isci.2021.103324.

Kolmogorov M, Armstrong J, Raney BJ, et al. 2018. Chromosome assembly of large and complex genomes using multiple references. Genome Res. 28(11):1720–1732. 10.1101/gr.236273.118.

Kolmogorov M, Raney B, Paten B, et al. 2014. Ragout—a reference-assisted assembly tool for bacterial genomes. Bioinformatics. 30(12):i302–i309. 10.1093/bioinformatics/btu280.

Krzywinski M, Schein J, Birol I, et al. 2009. Circos: an information aesthetic for comparative genomics. Genome Res. 19(9):1639–1645. 10.1101/gr.092759.109.

Kuang J-G, Han Z-T, Kang M-H, et al. 2023. Chromosome-level de novo genome assembly of two conifer-parasitic wasps, Megastigmus duclouxiana and Megastigmus sabinae, reveals genomic imprints of adaptation to hosts. Molecular Ecology Resources. 23(5):1142–1154. 10.1111/1755-0998.13785.

Kuznetsova VG, Golub NV. 2020. A checklist of chromosome numbers and a review of karyotype variation in Odonata of the world. Comp Cytogenet. 14(4):501–540. 10.3897/CompCytogen.v14i4.57062.

Kwak Y, Argandona JA, Degnan PH, et al. 2023. Chromosomal-level assembly of Bactericera cockerelli reveals rampant gene family expansions impacting genome structure, function and insect-microbe-plant-interactions. Molecular Ecology Resources. 23(1):233–252. 10.1111/1755-0998.13693.

L. S. Wolfe. 1952. A study of the genus Uropetala Selys (Order Odonata) from New Zealand. Transactions of the Royal Society of New Zealand. 80:245–275.

Laetsch DR, Blaxter ML. 2017. BlobTools: Interrogation of genome assemblies. F1000Res. 6:1287. 10.12688/f1000research.12232.1.

Lalejini A, Ferguson AJ, Grant NA, et al. 2021. Adaptive Phenotypic Plasticity Stabilizes Evolution in Fluctuating Environments. Frontiers in Ecology and Evolution. 9. Available from https://www.frontiersin.org/articles/10.3389/fevo.2021.715381.

Le Gal K, Schmidt EE, Sayin VI. 2021. Cellular Redox Homeostasis. Antioxidants (Basel). 10(9):1377. 10.3390/antiox10091377.

Li H. 2018. Minimap2: pairwise alignment for nucleotide sequences. Bioinformatics. 34(18):3094–3100. 10.1093/bioinformatics/bty191.

Li H. 2023. Protein-to-genome alignment with miniprot. Bioinformatics. 39(1):btad014. 10.1093/bioinformatics/btad014.

Lis C, Moore MP, Martin RA. 2020. Warm developmental temperatures induce non-adaptive plasticity in the intrasexually selected colouration of a dragonfly. Ecological Entomology. 45(3):663–670. 10.1111/een.12839.

Liu H, Jiang F, Wang S, et al. 2022. Chromosome-level genome of the globe skimmer dragonfly (Pantala flavescens). GigaScience. 11:giac009. 10.1093/gigascience/giac009.

Liu H-L, Harris AJ, Wang Z-F, et al. 2022. The genome of the Paleogene relic tree Bretschneidera sinensis: insights into trade-offs in gene family evolution, demographic history, and adaptive SNPs. DNA Research. 29(1):dsac003. 10.1093/dnares/dsac003.

Liu S, Hansen MM. 2017. PSMC (pairwise sequentially Markovian coalescent) analysis of RAD (restriction site associated DNA) sequencing data. Molecular Ecology Resources. 17(4):631–641. 10.1111/1755-0998.12606.

Luo S, Tang M, Frandsen PB, et al. 2018. The genome of an underwater architect, the caddisfly Stenopsyche tienmushanensis Hwang (Insecta: Trichoptera). GigaScience. 7(12):giy143. 10.1093/gigascience/giy143.

Manni M, Berkeley MR, Seppey M, et al. 2021. BUSCO Update: Novel and Streamlined Workflows along with Broader and Deeper Phylogenetic Coverage for Scoring of Eukaryotic, Prokaryotic, and Viral Genomes. Molecular Biology and Evolution. 38(10):4647–4654. 10.1093/molbev/msab199.

Marçais G, Kingsford C. 2011. A fast, lock-free approach for efficient parallel counting of occurrences of k-mers. Bioinformatics. 27(6):764–770. 10.1093/bioinformatics/btr011.

Mariani V, Biasini M, Barbato A, et al. 2013. lDDT: a local superposition-free score for comparing protein structures and models using distance difference tests. Bioinformatics. 29(21):2722–2728. 10.1093/bioinformatics/btt473.

May ML. 2013. A critical overview of progress in studies of migration of dragonflies (Odonata: Anisoptera), with emphasis on North America. J Insect Conserv. 17(1):1–15. 10.1007/s10841-012-9540-x.

May ML. 2019. Odonata: Who They Are and What They Have Done for Us Lately: Classification and Ecosystem Services of Dragonflies. Insects. 10(3):62. 10.3390/insects10030062.

Mendes FK, Vanderpool D, Fulton B, et al. 2021. CAFE 5 models variation in evolutionary rates among gene families. Bioinformatics. 36(22–23):5516–5518. 10.1093/bioinformatics/btaa1022.

Misof B, Liu S, Meusemann K, et al. 2014. Phylogenomics resolves the timing and pattern of insect evolution. Science. 346(6210):763–767. 10.1126/science.1257570

Newton L, Tolman E, Kohli M, et al. 2023 Jun 7. Evolution of Odonata: Genomic insights. Current Opinion in Insect Science.:101073. 10.1016/j.cois.2023.101073.

Parkes K, Amos W, Moore N, et al. 2009. Population structure and speciation in the dragonfly Sympetrum striolatum/nigrescens (Odonata: Libellulidae): An analysis using AFLP markers. European Journal of Entomology. 106:179–184. 10.14411/eje.2009.021.

F. Petrulevičius J, Nel A. 2003. Oldest Petalurid dragonfly (Insecta: Odonata): a Lower Cretaceous specimen from south Patagonia, Argentina. Cretaceous Research. 24(1):31–34. 10.1016/S0195-6671(03)00025-9

Phillips L. 2012. gnuplot Cookbook. Birmingham: Packt Publishing. 220 p.

Price BW, Winter M, Brooks SJ. 2022. The genome sequence of the blue-tailed damselfly, Ischnura elegans (Vander Linden, 1820). Wellcome Open Res. 7:66. 10.12688/wellcomeopenres.17691.1.

R F, R K-M, M K, et al. 2015. Extraordinary diversity of visual opsin genes in dragonflies. Proceedings of the National Academy of Sciences of the United States of America. 112(11). 10.1073/pnas.1424670112.

Ranallo-Benavidez TR, Jaron KS, Schatz MC. 2020. GenomeScope 2.0 and Smudgeplot for reference-free profiling of polyploid genomes. Nat Commun. 11(1):1432. 10.1038/s41467-020-14998-3.

Ranson H, Claudianos C, Ortelli F, et al. 2002. Evolution of supergene families associated with insecticide resistance. Science. 298(5591):179–181. 10.1126/science.1076781

Rebora M, Salerno G, Piersanti S. 2022. Odonata perception is more than vision. In: Cordoba-Aguilar A, Beatty C, Bried J, editors. Dragonflies and Damselflies: Model Organisms for Ecological and Evolutionary Research. Oxford University Press. p. 0. 10.1093/oso/9780192898623.003.0007.

Rowe RJ. 1981. AN ANNOTATED KEY TO THE NEW ZEALAND ODONATA. 9.

Rowe RJ. 2004. Conservation of Odonata in the South Pacific and Australasia. International Journal of Odonatology. 7(2):139–147. 10.1080/13887890.2004.9748206.

Russell RW, May ML, Soltesz KL, et al. 1998. Massive Swarm Migrations of Dragonflies (Odonata) in Eastern North America. amid. 140(2):325–342. 10.1674/0003-0031(1998)140[0325:MSMODO]2.0.CO;2.

Salmón P, Watson H, Nord A, et al. 2018. Effects of the Urban Environment on Oxidative Stress in Early Life: Insights from a Cross-fostering Experiment. Integr Comp Biol. 58(5):986–994. 10.1093/icb/icy099.

Sharip Z, Fauzi M. 2019. Microbial Contamination in Urban Tropical Lentic Waterbodies and Ponds along Urbanization Gradient. Pertanika Journal of Tropical Agricultural Science. 42.

Shingate P, Ravi V, Prasad A, et al. 2020. Chromosome-level assembly of the horseshoe crab genome provides insights into its genome evolution. Nature Communications. 11(1):2322. 10.1038/s41467-020-16180-1.

Shiryev SA, Papadopoulos JS, Schäffer AA, et al. 2007. Improved BLAST searches using longer words for protein seeding. Bioinformatics. 23(21):2949–2951. 10.1093/bioinformatics/btm479.

Šigutová H, Dolný A, Samways MJ, et al. 2022. Odonata as indicators of pollution, habitat quality, and landscape disturbance. In: Cordoba-Aguilar A, Beatty C, Bried J, editors. Dragonflies and Damselflies: Model Organisms for Ecological and Evolutionary Research. Oxford University Press. p. 0. 10.1093/oso/9780192898623.003.0026.

[Software] R Core Team. 2021. R: A language and environment for statistical computing.

Stanke M, Tzvetkova A, Morgenstern B. 2006. AUGUSTUS at EGASP: using EST, protein and genomic alignments for improved gene prediction in the human genome. Genome Biology. 7(1):S11. 10.1186/gb-2006-7-s1-s11.

Supek F, Bošnjak M, Škunca N, et al. 2011. REVIGO Summarizes and Visualizes Long Lists of Gene Ontology Terms. Gibas C, editor. PLoS ONE. 6(7):e21800. 10.1371/journal.pone.0021800.

Suvorov A, Jensen NO, Sharkey CR, et al. 2017. Opsins have evolved under the permanent heterozygote model: insights from phylotranscriptomics of Odonata. Molecular Ecology. 26(5):1306–1322. 10.1111/mec.13884.

Suvorov A, Scornavacca C, Fujimoto MS, et al. 2021 Jul 29. Deep Ancestral Introgression Shapes Evolutionary History of Dragonflies and Damselflies. Systematic Biology.:syab063. 10.1093/sysbio/syab063.

Teets NM, Peyton JT, Colinet H, et al. 2012. Gene expression changes governing extreme dehydration tolerance in an Antarctic insect. Proceedings of the National Academy of Sciences. 109(50):20744–20749. 10.1073/pnas.1218661109.

Tennessen K. 2019. Libellulidae. In: Tennessen KJ, editor. Dragonfly Nymphs of North America: An Identification Guide. Cham: Springer International Publishing. p. 407–576. 10.1007/978-3-319-97776-8_12.

Thomas GWC, Dohmen E, Hughes DST, et al. 2020. Gene content evolution in the arthropods. Genome Biology. 21(1):15. 10.1186/s13059-019-1925-7

Tollett VD, Benvenutti EL, Deer LA, et al. 2009. Differential Toxicity to Cd, Pb, and Cu in Dragonfly Larvae (Insecta: Odonata). Arch Environ Contam Toxicol. 56(1):77–84. 10.1007/s00244-008-9170-1.

Tolman ER, Beatty CD, Bush J, et al. 2023. Exploring chromosome evolution in 250 million year old groups of dragonflies and damselflies (Insecta:Odonata). Mol Ecol. 32(21):5785–5797. 10.1111/mec.17147.

Tolman ER, Beatty CD, Bush J, et al. 2023 Feb 18. A Chromosome-length Assembly of the Black Petaltail (Tanypteryx hageni) Dragonfly. Genome Biology and Evolution.:evad024. 10.1093/gbe/evad024.

Tolman ER, Beatty CD, Kohli MK, et al. 2024. A molecular phylogeny of the Petaluridae (Odonata: Anisoptera): A 160-Million-Year-Old story of drift and extinction. Molecular Phylogenetics and Evolution. 200:108185. 10.1016/j.ympev.2024.108185.

Tolman ER, Gamett E, Beatty CD, et al. 2024. The Blueprint for Survival: The Blue Dasher Dragonfly as a Model for Urban Adaptation. :2024.12.12.628234. 10.1101/2024.12.12.628234

Tolman ER, Bruchim OR, Driever ES, et al. 2023. Changes in effective population size of Odonata in response to climate change revealed through genomics. IJO. 26:205–211. 10.48156/1388.2023.1917241.

Trizna M. 2020. assembly_stats 0.1.4. 10.5281/zenodo.3968775.

Turner PE. 1970. Allusive Dragons: Functional Integration and Structural Integrity among Natural Populations of the Relict Dragonfly Tanyptery hageni. [PhD Thesis]. Berkeley, California: University of California, Berkeley.

Uche-Dike R, Tolman ER, Benischek C, et al. 2024. Environmental DNA vs. Community Science: Strengths and Limitations for Urban Odonata Surveys. :2024.11.26.625270. 10.1101/2024.11.26.625270

Uliano-Silva M, Nunes JGF, Krasheninnikova K, et al. 2021. marcelauliano/MitoHiFi: mitohifi_v2.0. 10.5281/zenodo.5205678.

Vollmer W, Blanot D, De Pedro MA. 2008. Peptidoglycan structure and architecture. FEMS Microbiology Reviews. 32(2):149–167. 10.1111/j.1574-6976.2007.00094.x.

Waller JT, Willink B, Tschol M, et al. 2019. The odonate phenotypic database, a new open data resource for comparative studies of an old insect order. Sci Data. 6(1):316. 10.1038/s41597-019-0318-9

Wang Y, Luan Y, Luo J, et al. 2023. 300 Million years of coral treaders (Insecta: Heteroptera: Hermatobatidae) back to the ocean in the phylogenetic context of Arthropoda. Proceedings of the Royal Society B: Biological Sciences. 290(2001):20230855. 10.1098/rspb.2023.0855.

Wang Y, Tang H, DeBarry JD, et al. 2012. MCScanX: a toolkit for detection and evolutionary analysis of gene synteny and collinearity. Nucleic Acids Res. 40(7):e49. 10.1093/nar/gkr1293.

Ware J, Kohli MK, Mendoza CM, et al. 2022. Evidence for widespread gene flow and migration in the Globe Skimmer dragonfly Pantala flavescens. International Journal of Odonatology. 25:43–55.

Ware JL, Beatty CD, Herrera MS, et al. 2014a. The petaltail dragonflies (Odonata: Petaluridae): Mesozoic habitat specialists that survive to the modern day. Journal of Biogeography. 41(7):1291–1300. 10.1111/jbi.12273.

Ware JL, Beatty CD, Herrera MS, et al. 2014b. The petaltail dragonflies (Odonata: Petaluridae): Mesozoic habitat specialists that survive to the modern day. Journal of Biogeography. 41(7):1291–1300. 10.1111/jbi.12273.

Winstanley WJ, Rowe RJ. 1980. The larval habitat of Uropetala carovei carovei (Odonata: Petaluridae) in the North Island of New Zealand, and the geographical limits of the subspecies. New Zealand Journal of Zoology. 7(1):127–134. 10.1080/03014223.1980.10423769.

Wissinger SA. 1988. Life History and Size Structure of Larval Dragonfly Populations. Journal of the North American Benthological Society. 7(1):13–28. 10.2307/1467827.

Zhang J, Xie P, Lascoux M, et al. 2013. Rapidly Evolving Genes and Stress Adaptation of Two Desert Poplars, Populus euphratica and P. pruinosa. PLOS ONE. 8(6):e66370. 10.1371/journal.pone.0066370

