## supplementary materials for "Newly Sequenced Genomes Reveal Patterns of Gene Family Expansion in Select Dragonflies (Odonata: Anisoptera)"

Supplementary figure 1: *Genomescope profiles of Pachydiplax longipennis and Pantala flavescens*

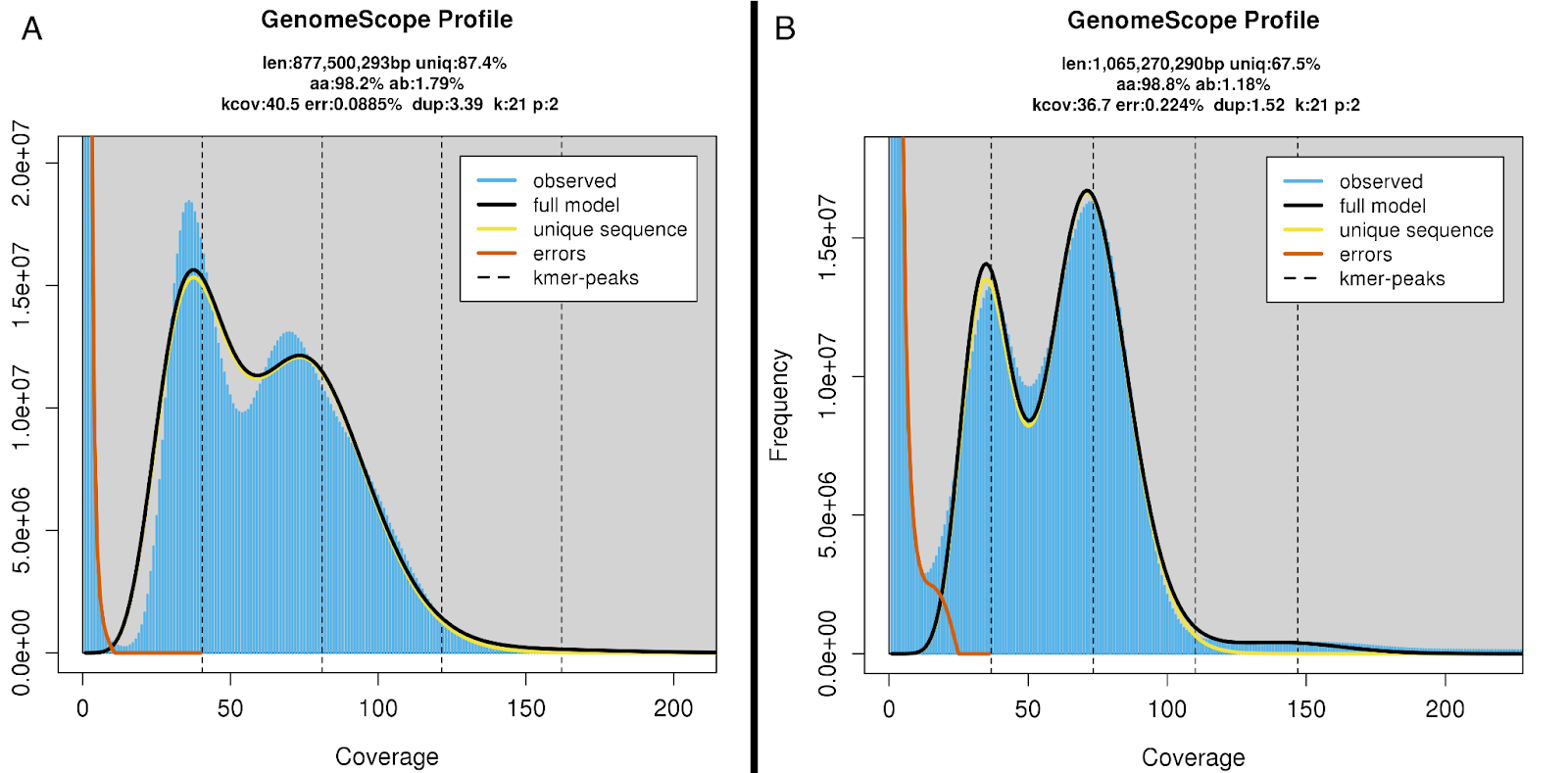

*Supplementary figure 1: Genomescope (Ranallo-Benavidez et al., 2020) profile of (A) Pachydiplax longipennis and (B) P. flavescens.*

*Supplementary figure 2: Blobplots of Uropetala carovei and P. longipennis*

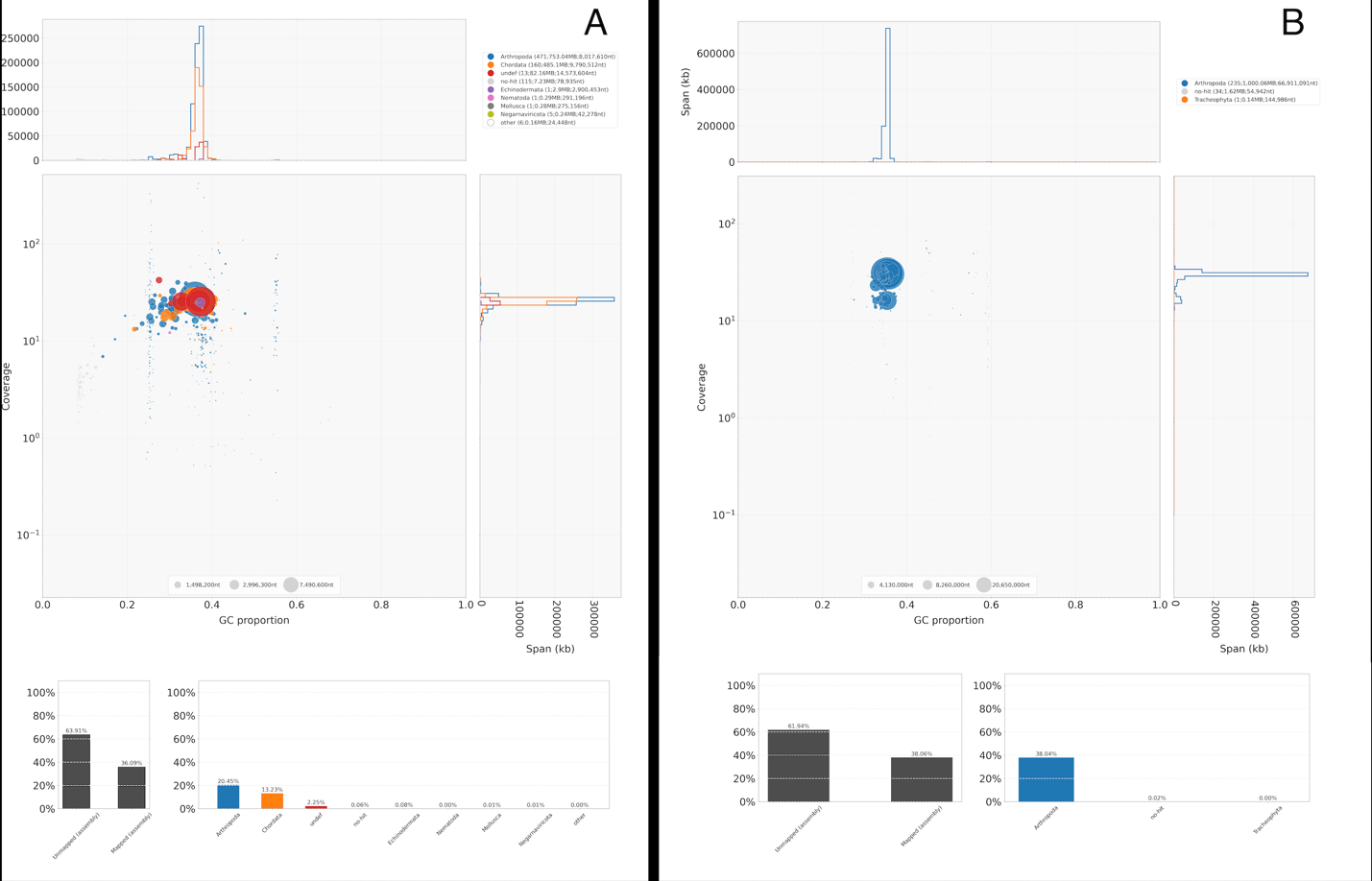

*Supplementary figure 2: Bloplots (Laetsch & Blaxter, 2017) of (A) Uropetala carovei and (B) Pachydiplax longipennis.*

Supplementary figure 3: *Synteny between the genome assemblies of S. striolatum and P. longipennis*

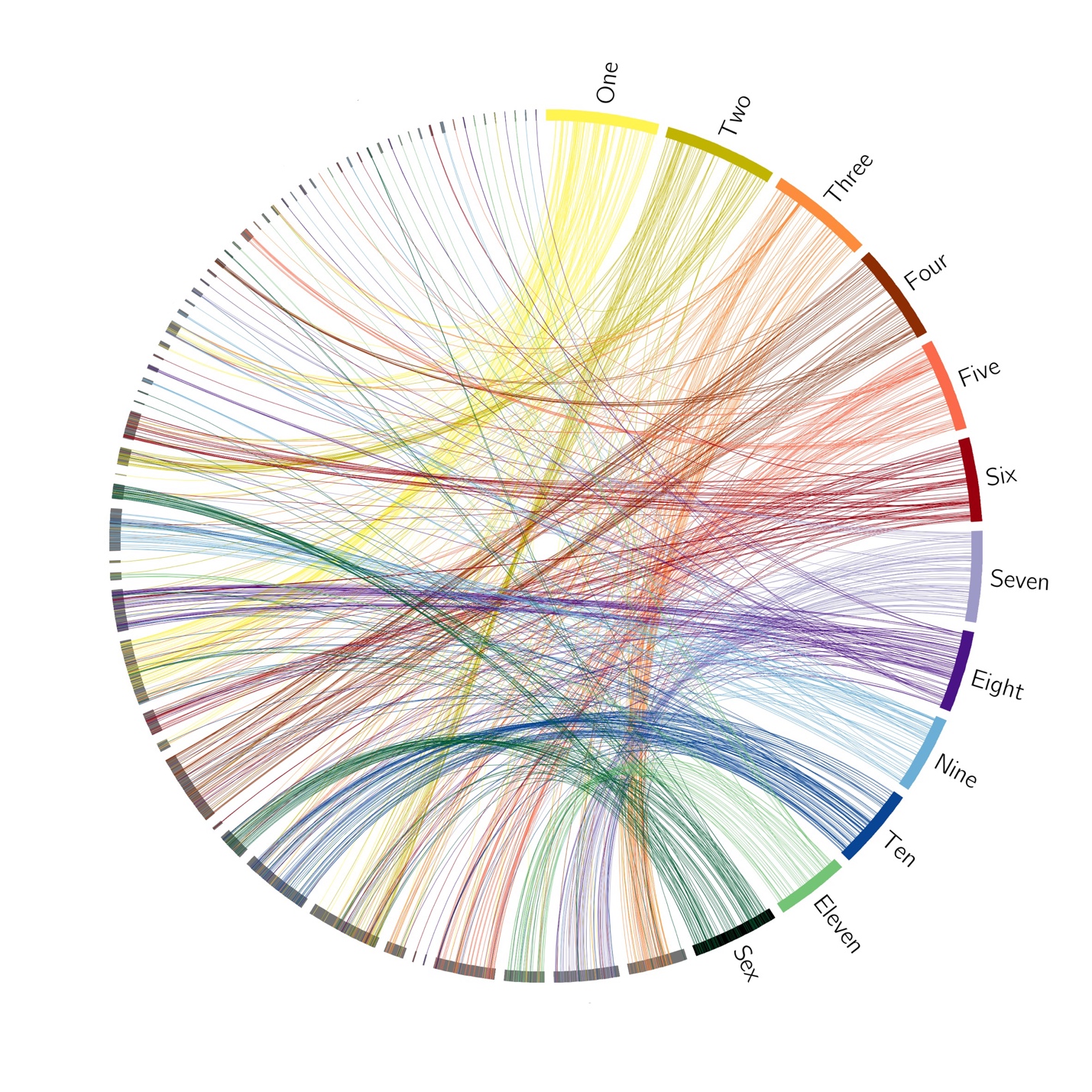

*Supplementary figure 3: Circos (Krzywinski et al., 2009) plot showing synteny between the chromosome-length genome assembly of S. striolatum (Crowley et al., 2023) and the draft assembly of P. longipennis. Contigs from the assembly of P. longipennis are shown in grey.*

Supplementary figure 4: *Density plots of -LnL and Global Lambda Estimates from 1000 Global Lambda Model Runs*
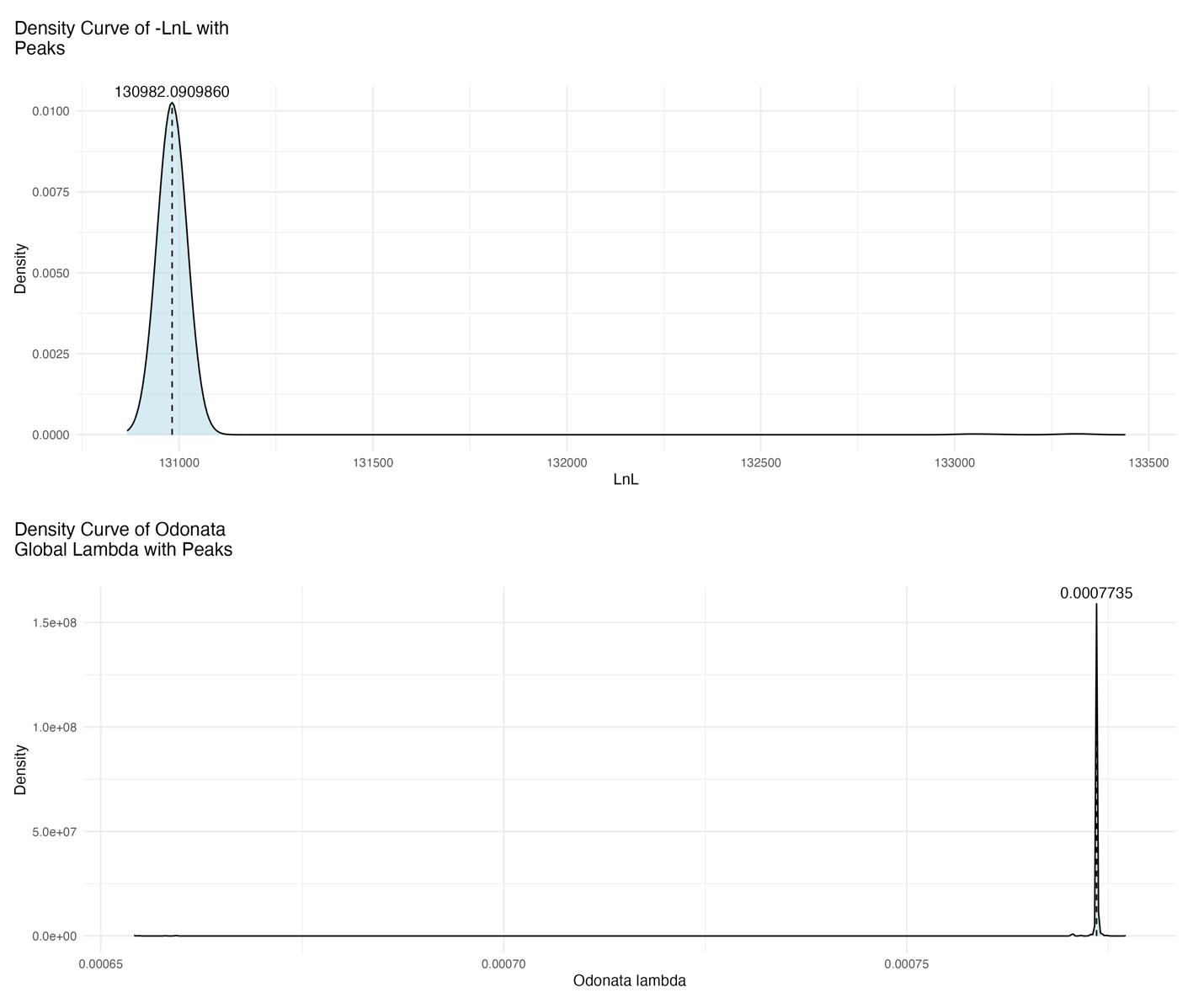

*Supplementary figure 4: Density of model -LnL (top) and the global lambda estimates (bottom) from 1,000 independent CAFE5 runs of a global lambda model.*

Supplementary figure 5: *Density plots of -LnL and Lambda Estimates from 1000 Two Lambda Model Runs*
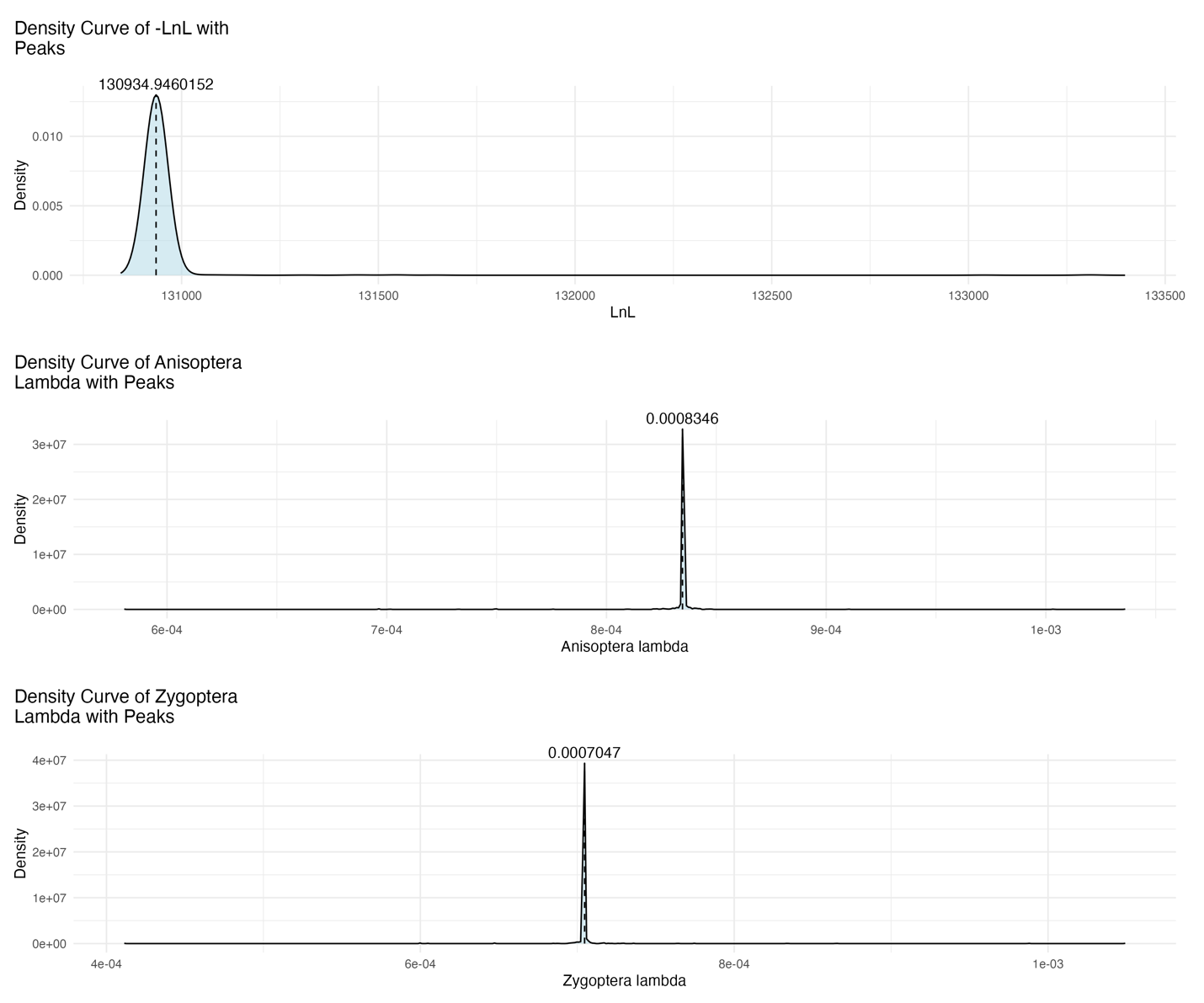

*Supplementary figure 5: Density of model -LnL (top), Anisoptera lambda estimates (middle) and Zygoptera lambda estimates (bottom) from 1,000 independent CAFE5 runs of a two lambda model.*

Supplementary figure 6: *Density plots of -LnL and Lambda Estimates from 1000 Four Lambda Model Runs*

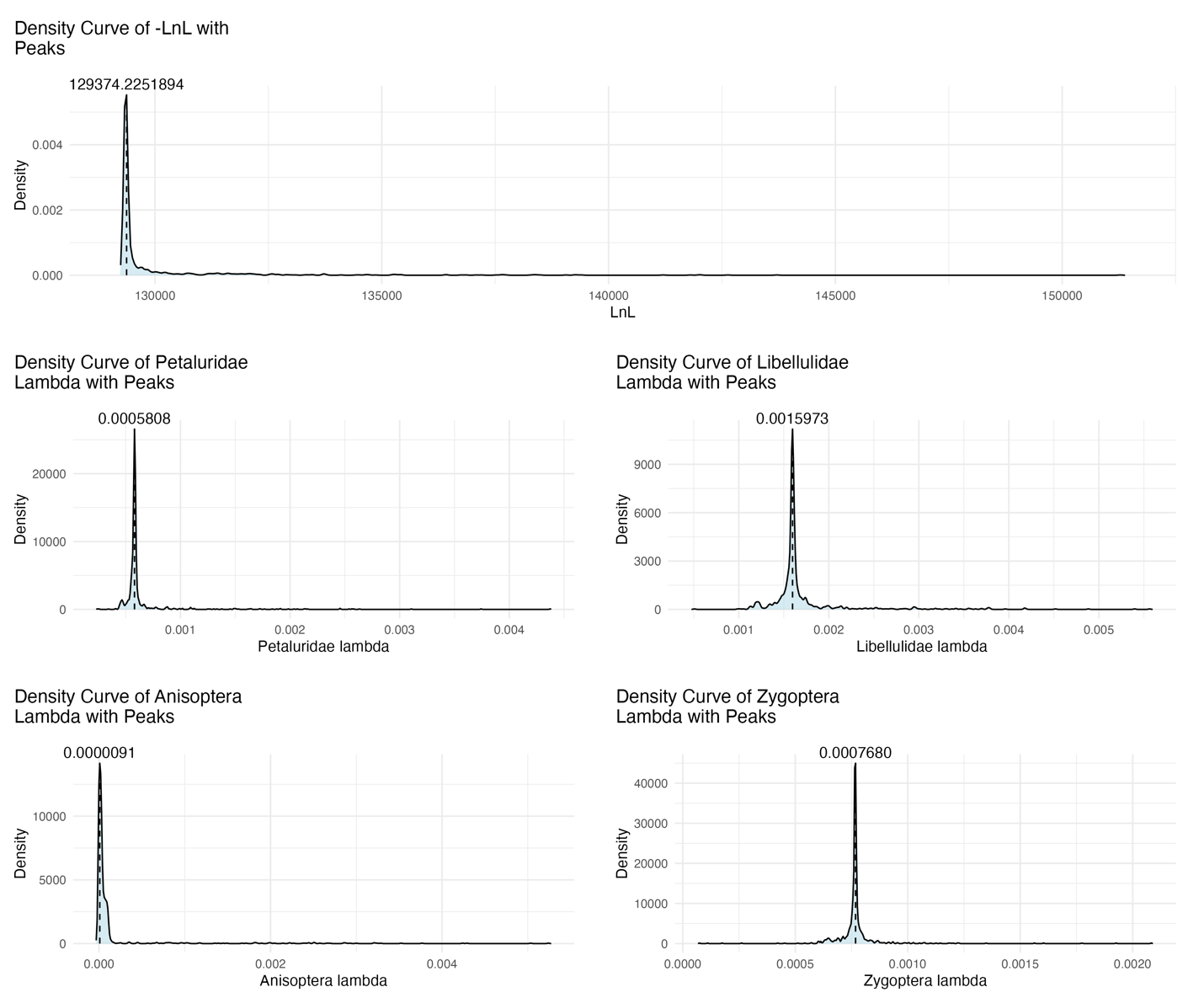

*Supplementary figure 6: Density of model -LnL (top), Petaluridae and Libellulidae lambda estimates (middle), and Anisoptera and Zygopyera lambda estimates (bottom) from 1,000 independent CAFE5 runs of a four lambda model.*

| Supplementary Table 1: Contaminants removed from *Uropetala carovei* assembly with NCBI contamination filter | | |
| --- | --- | --- |
| Contig | Size | Apparent Source |
| ptg000435l | 92003 | anml:primates |
| ptg000438l | 31341 | anml:primates |
| ptg000471l | 17420 | prok:mycoplasmas |
| ptg000480l | 30935 | anml:primates |
| ptg000500l | 42100 | anml:primates |
| ptg000504l | 36509 | anml:primates |
| ptg000505l | 35933 | anml:primates |
| ptg000516l | 31804 | anml:primates |
| ptg000539l | 28657 | anml:primates |
| ptg000565l | 62872 | anml:primates |
| ptg000571l | 33519 | prok:b-proteobacteria |
| ptg000626l | 28876 | anml:primates |
| ptg000635l | 84022 | anml:primates |
| ptg000643l | 32577 | anml:primates |
| ptg000672l | 42278 | anml:primates |
| ptg000689l | 40170 | anml:primates |
| ptg000697l | 24224 | prok:b-proteobacteria |
| ptg000700l | 31912 | anml:primates |
| ptg000715l | 20572 | prok:b-proteobacteria |
| ptg000727l | 27495 | anml:primates |
| ptg000735l | 32693 | anml:primates |
| ptg000738l | 28200 | anml:primates |
| ptg000744l | 27256 | anml:primates |
| *Supplementary table 1: Contaminants removed from the assembly of U. carovei with the NCBI contamination filter.* | | |

| Supplementary Table 2: Synteny between the genomes of *P. longipennis* and *S. striolatum* [[31]](https://www.zotero.org/google-docs/?Hkylly) | | |
| --- | --- | --- |
| Chromosome (from *Sympetrum striolatum* [[31]](https://www.zotero.org/google-docs/?kZUlG2)*)* | Syntenic contigs from *P. longipennis* | Contigs used in scaffolding |
| One | ptg000002l(2), ptg000003l(1), ptg000004l(3), ptg000007l(2), ptg000008l(6), ptg000009l(2), ptg000010l(1), ptg000012l(3), ptg000013l(4), ptg000015l(54), ptg000016l(3), ptg000019l(1), ptg000020l(2), ptg000022l(1), ptg000024l(2), ptg000026l(1), ptg000032l(2), ptg000034l(7), ptg000040l(1), ptg000050l(4) | Ptg000053l, ptg000013l, ptg000050l , ptg000015l*, ptg000034l*, ptg000032l* |
| Two | ptg000001l(4), ptg000002l(1), ptg000003l(1), ptg000004l(2), ptg000007l(1), ptg000008l(26), ptg000009l(5), ptg000012l(2), ptg000014l(1), ptg000015l(4), ptg000016l(4), ptg000017l(1), ptg000018l(1), ptg000020l(1), ptg000024l(17), ptg000026l(3), ptg000034l(1), ptg000095l(1 | ptg000008l , ptg000024l |
| Three | ptg000001l(38), ptg000002l(4), ptg000003l(3), ptg000007l(10), ptg000008l(4), ptg000009l(5), ptg000010l(3), ptg000012l(1), ptg000014l(2), ptg000015l(1), ptg000016l(1), ptg000019l(2), ptg000020l(1), ptg000026l(2), ptg000034l(4), ptg000047l(1), ptg000050l(3), ptg000058l(1) | Ptg000047l, ptg000001l , ptg000007l |
| Four | ptg000001l(1), ptg000002l(2), ptg000004l(1), ptg000010l(2), ptg000012l(52), ptg000040l(8) | - |
| Five | ptg000001l(3), ptg000003l(1), ptg000004l(37), ptg000008l(6), ptg000009l(5), ptg000010l(1), ptg000012l(5), ptg000013l(1), ptg000014l(2), ptg000015l(9), ptg000034l(5), ptg000044l(9), ptg000058l(1), ptg000089l(1) | - |
| Six | ptg000002l(2), ptg000004l(3), ptg000006l(1), ptg000008l(4), ptg000009l(2), ptg000010l(3), ptg000011l(3), ptg000012l(4), ptg000014l(16), ptg000015l(1), ptg000016l(4), ptg000019l(1), ptg000020l(1), ptg000024l(1), ptg000026l(19), ptg000031l(1), ptg000039l(1), ptg000063l(1), ptg000066l(1), ptg000085l(2) | Ptg000026l, ptg000014l |
| Seven | ptg000001l(2), ptg000002l(42), ptg000003l(2), ptg000004l(3), ptg000007l(1), ptg000008l(2), ptg000009l(3), ptg000010l(1), ptg000015l(1), ptg000016l(1), ptg000019l(1), ptg000020l(1), ptg000024l(1), ptg000034l(3), ptg000037l(4), ptg000040l(1), ptg000050l(1) | - |
| Eight | ptg000002l(4), ptg000004l(1), ptg000005l(1), ptg000007l(1), ptg000008l(4), ptg000009l(5), ptg000010l(2), ptg000012l(3), ptg000014l(2), ptg000016l(29), ptg000024l(1), ptg000030l(4), ptg000034l(1), ptg000038l(1), ptg000040l(1), ptg000051l(2), ptg000054l(1), ptg000075l(2), ptg000090l(2), ptg000098l(1), ptg000103l(1) | ptg000038l, ptg000103l |
| Nine | ptg000002l(1), ptg000003l(1), ptg000009l(3), ptg000014l(1), ptg000015l(3), ptg000019l(25), ptg000029l(4), ptg000034l(5), ptg000035l(3), ptg000036l(1), ptg000052l(1), ptg000056l(1), ptg000065l(2), ptg000083l(1), ptg000086l(2), ptg000100l(1) | ptg000019l , ptg000035l |
| Ten | ptg000001l(1), ptg000002l(2), ptg000003l(1), ptg000004l(6), ptg000008l(5), ptg000009l(51), ptg000010l(3), ptg000012l(4), ptg000013l(1), ptg000014l(1), ptg000015l(6), ptg000016l(5), ptg000019l(4), ptg000020l(2), ptg000024l(1), ptg000037l(1) | - |
| Eleven | ptg000003l(25), ptg000009l(3), ptg000010l(1), ptg000015l(2), ptg000017l(4), ptg000020l(1), ptg000042l(2), ptg000049l(1), ptg000050l(1), ptg000060l(2), ptg000074l(1), ptg000077l(1), ptg000081l(1), ptg000091l(2), ptg000093l(1), ptg000099l(2) | - |
| Sex | ptg000001l(4), ptg000002l(2), ptg000003l(2), ptg000007l(1), ptg000008l(6), ptg000009l(2), ptg000010l(28), ptg000015l(4), ptg000016l(2), ptg000019l(1), ptg000020l(20), ptg000024l(1), ptg000026l(1), ptg000027l(1), ptg000028l(1), ptg000040l(1), ptg000041l(3), ptg000068l(3) | Ptg000010l, ptg000020l |
| *Supplementary table 2: Lists contigs of P. longipennis which have synteny with chromosomes of S. striolatum* (Crowley et al., 2023)*, and contigs scaffolded into pseudo chromosomes.*  *Unlocalized | | |

| Lambda model | Converged -LnL | Parameters | Converged model AIC | Percentage of 1000 iterations model is chosen by AIC | Converged model LRT | Percentage of 1000 iterations model is chosen by LRT | SD of -LnL from 1000 independent runs |
| --- | --- | --- | --- | --- | --- | --- | --- |
| Global lambda | 130982 | 1 | 261966 | 0.3% | - | 0.3% | 170.67 |
| Separate lambdas for Anisoptera and Zygoptera | 130935 | 2 | 261974 | 11.3% | 94** | 11.3% | 132.67** |
| Separate lambdas for ancestral Anisoptera, Petaluridae, Libellulidae and Zygoptera | 129347 | 4 | 258702 | 88.4% | 3270** | 88.4% | 1872.92** |
| Supplementary table three: Comparison of Lambda model fit and convergence  ** significantly differs from global lambda model p < 0.001 | | | | | | | |

**References**

Crowley, L. M., Price, B. W., Allan, E. L., Eagles, M., University of Oxford and Wytham Woods Genome Acquisition Lab, Natural History Museum Genome Acquisition Lab, Darwin Tree of Life Barcoding collective, Wellcome Sanger Institute Tree of Life programme, Wellcome Sanger Institute Scientific Operations: DNA Pipelines collective, Tree of Life Core Informatics collective, & Darwin Tree of Life Consortium. (2023). The genome sequence of the Common Darter, Sympetrum striolatum (Charpentier, 1840). *Wellcome Open Research*, *8*, 389. https://doi.org/10.12688/wellcomeopenres.19937.1

Krzywinski, M., Schein, J., Birol, I., Connors, J., Gascoyne, R., Horsman, D., Jones, S. J., & Marra, M. A. (2009). Circos: An information aesthetic for comparative genomics. *Genome Research*, *19*(9), 1639–1645. https://doi.org/10.1101/gr.092759.109

Laetsch, D. R., & Blaxter, M. L. (2017). BlobTools: Interrogation of genome assemblies. *F1000Research*, *6*, 1287. https://doi.org/10.12688/f1000research.12232.1

Mendes, F. K., Vanderpool, D., Fulton, B., & Hahn, M. W. (2021). CAFE 5 models variation in evolutionary rates among gene families. *Bioinformatics*, *36*(22–23), 5516–5518. https://doi.org/10.1093/bioinformatics/btaa1022

Ranallo-Benavidez, T. R., Jaron, K. S., & Schatz, M. C. (2020). GenomeScope 2.0 and Smudgeplot for reference-free profiling of polyploid genomes. *Nature Communications*, *11*(1), Article 1. https://doi.org/10.1038/s41467-020-14998-3
